# PKA drives paracrine crisis and WNT4-dependent testis tumor in Carney complex

**DOI:** 10.1101/2020.12.21.423735

**Authors:** C. Djari, I. Sahut-Barnola, A. Septier, I. Plotton, N. Montanier, D. Dufour, A. Levasseur, J. Wilmouth, JC. Pointud, FR. Faucz, C. Kamilaris, AG. Lopez, F. Guillou, A. Swain, S. Vainio, I. Tauveron, P. Val, H. Lefebvre, CA. Stratakis, A. Martinez, AM. Lefrançois-Martinez

## Abstract

Large Cell Calcifying Sertoli Cell Tumors (LCCSCTs) are among the most frequent lesions occurring in Carney complex (CNC) male patients. Although they constitute a key diagnostic criterion for this rare multiple neoplasia syndrome resulting from inactivating mutations of the tumor suppressor *PRKAR1A* leading to unrepressed PKA activity, the LCCSCT pathogenesis and origin remain elusive. Mouse models targeting *Prkar1a* inactivation in all somatic populations or separately in each cell type were generated to decipher the molecular and paracrine networks involved in the CNC testis lesion induction. We demonstrate that *Prkar1a* mutation is required in both stromal and Sertoli cells for the occurrence of LCCSCT. Integrative analyses comparing transcriptomic, immunohistological data and phenotype of mutant mouse combinations led to understand the human LCCSCT pathogenesis and demonstrated unprecedented PKA-induced paracrine molecular circuits in which the aberrant WNT4 signal production is a limiting step in shaping intratubular lesions and tumor expansion both in mouse model and human CNC testes.

## INTRODUCTION

Carney complex (CNC) is a rare multiple neoplasia syndrome mainly associated with inactivating mutation of *PRKAR1A* gene encoding the R1α subunit of the PKA which results in overactive PKA signaling pathway (1). It is characterized by pigmented lesions of the skin and mucosae, cardiac, cutaneous and other myxomatous tumors, and multiple other endocrine and non-endocrine neoplasms (2, 3). Approximatively 70% of patients are estimated to develop Large Cell Calcifying Sertoli Cell Tumors (LCCSCTs) (2, 4). LCCSCT is a testicular Sex Cord Stromal Tumor (SCST) subtype characterized by bilateral and multifocal masses caused by inactivating mutations of *PRKAR1A* (17q22-24) associated with Carney complex (CNC) but is also described in Peutz-Jeghers syndrome (PJS) caused by inactivating mutations of *STK11* (19p13.3) (5, 6). LCCSCTs developing in the case of CNC (CNC-LCCSCTs) are mostly benign tumors with low malignant potential and may be diagnosed from the childhood by sonographic examination, and associated with precocious puberty in some cases (7, 8). In engineered *Prkar1a^+/−^* mice, loss of heterozygosity occurrence for *Prkar1a* locus has been implicated in the tumorigenesis process for CNC related neoplasia (9) indicating that the resulting constitutive PKA activity plays an oncogenic role in specific tissues (10). The LCCSCT current management consists in the resection of the primary tumor owing to the limited effect of chemotherapy or radiation treatments. However the tumor discriminant markers are insufficient to allow early detection of a potential recurrence and to distinguish LCCSCT from other SCST. Histological analyses highlight various tissue alterations (11). Moderate tubular damages characterized by spermatogenesis defects, calcifications, peritubular fibrosis and composite tumour mass combined with Sertoli cell proliferating nodules are described (12, 13). LCCSCT are systematically associated with increased production of Inhibin-α, S100, Calretinin and β-catenin but its molecular status is still incomplete due to the low LCCSCT incidence (11, 14, 15). Therefore the molecular mechanisms and the cellular events involved in LCCSCT pathogenesis remain to be understood to improve their management. Moreover, the reproductive alterations have been poorly investigated in male CNC patients, concerning their fertility as well as their testis endocrine activity.

In embryo, Sertoli cells (SC) and Leydig cells (LC) arise from a common pool of SF1+ (also called NR5A1) progenitor cells under the control of coordinate genetic and paracrine mechanisms triggered by the male determining SRY and SOX9 induced programs to counteract the female RSPO1/WNT4/β-catenin and FOXL2 pathways (16). At adulthood, testicular physiology carries out exocrine and endocrine functions: SC support the germ cell (GC) nursing including growth and differentiation in seminiferous tubules (ST), and LC produce androgens in interstitial compartment. Pituitary Follicle Stimulating Hormone (FSH) and Luteinizing Hormone (LH) are the main regulators of the adult SC and LC functions through their respective specific G protein-coupled receptor, FSH-R and LH-R, both interacting with number of intracellular effectors, among which the cAMP/PKA pathway plays a central role. Activating mutations of human FSH-R and LH-R resulting in permanent PKA activity have been described for several years and their physiological outcomes were extensively studied through biochemical and functional approaches including mutant mice models (17–20). Whether activating mutations of the gonadotropin receptors are involved in tumor development is not yet clear and their pathological outcomes do not overlap with the testis CNC lesions etiology. The objective of our work was to identify the molecular networks (deregulated by PKA activation) which cause the resulting cellular alterations found in the CNC testis. We generated several mouse genetic models lacking *Prkar1a* either in all somatic testicular cells arising from fetal somatic progenitor (prKO model: *Sf1:Cre,Prkar1a^fl/fl^*), or exclusively in SC (srKO model: *Amh:Cre,Prkar1a^fl/fl^*) or LC (lrKO model: *Cyp11a1:Cre,Prkar1a^fl/fl^*). The ontogenic follow-up of these models together with genetic manipulation of catalytic PKA subunit and WNT signal should provide a readout of the sequence of histological, molecular and endocrine modifications arising in human LCCSCT from CNC patient.

## RESULTS

### *Prkar1a* loss in mouse testicular somatic cells induces LCCSCT-like lesions

*Sf1:Cre, Prkar1a^fl/fl^* mice (prKO mice) were generated to clarify the CNC-LCCSCT lesions arising mechanisms by targeting early *Prkar1a* inactivation (E10.5) in somatic testis cells (Figure S1A and B). The use of *Rosa26R-mtmg* reporter locus confirmed that the recombination affected all somatic testis cell type excluding the GC population as previously described (21) (Figure S1 A-C) and RTqPCR analyses confirmed a robust reduction in *Prkar1a* mRNA levels in prKO testes (Figure S1D). The mutant mice colony setup revealed that prKO male mice were sterile since the onset of puberty due to a lack of spermatozoa production and displayed disrupted gonadotropic axis with collapsed FSH and LH levels associated with hypertestosteronemia (Figure S1 E-G). By contrast, 2-month-old *Sf1:cre,Prkar1a^fl/fl^,Prkaca^−/+^* male mice, harboring *Prkaca* haploinsufficiency with decreased PKA catalytic activity, retained healthy testis histological integrity indicating that testis failures in prKO mice were mostly due to PKA overactivation (Figure S2 A-E). Masson trichrome and HE staining revealed that 2 and 4-month-old prKO mice exhibited a severe gonadic dysplasia with seminiferous compartment disorganization. These defects were combined to calcification areas revealed by Alizarin Red coloration, peritubular basal lamina thickening attested by Laminin B1 immunostaining, and germ cell loss in close similarity with human CNC-LCCSCT histology (Figure 1, A and B) (Figure S1 H-J) (Figure S3). Transcripts levels for the SCST current molecular markers *i.e. Calb2, Ctnnb1, Inha, S100a1* were also strongly increased in the 4-month-old prKO testes compared with WT (Figure 1C). Moreover, hypertrophic blood vessels and early bilateral proliferative stromal tumor mass expansion attested by increased stromal PCNA index were observed in 4-month-old mutant testes whereas only limited stromal hyperplastic areas arose without tubular defect in 2-week-old mutant testis (Figure 1A, D-E) (Figure S1K-L). By contrast, heterozygous *Sf1:Cre*,*Prkar1a^fl/+^* littermates display healthy testicular phenotype similarly to the *Prkar1a^fl/fl^* mice. All these observations demonstrated that loss of *Prkar1a* in mice phenocopied all tissue defects found in testes from CNC patients. To gain insight into the gene expression profiles associated with the prKO testis lesion set up, we conducted comparative microarray analyses on gene expression in 2-week and 2-month-old prKO and WT testes. Transcriptomic analyses revealed 1328 significantly deregulated genes in adult prKO mice (676 up- and 652 down-regulated genes) whereas only 93 genes were deregulated at 2 weeks in mutant testes (90 up- and 3 down-regulated genes compared with WT (abs(Log2FC)>1, adj. p-val.<0.05) (Figure 1F and G). These data emphasized the delayed functional response to *Prkar1a* inactivation that was triggered from the early fetal gonadal stage (Figure S1B). Gene ontology analyses performed on transcriptomic data from 2-week-old testes highlighted an increased expression in genes involved in steroid metabolism whereas gene signatures associated with intratubular cells homeostasis were virtually unaffected (Datafiles S3). Consistent with these observations, hormonal assays revealed a significative 134-fold increased testosterone levels in 17 day-old prKO testes (5.39ng/organ) compared with WT (0.04ng/organ) (plasma testosterone levels = 16.01ng/mL ± 5.12 in prKO vs 0.16 ng/mL ± 0.04 in WT) whereas testosterone levels were unchanged in the plasma of 5-day-old prKO mice (0.27ng/mL ± 0.12) compared with WT (0.14ng/mL ± 0.04). In 2-month-old prKO mice, unbiased Gene Ontology (GO) and KEGG analyses revealed a negative enrichment in molecular signatures associated with spermatogenesis, sperm motility and a positive enrichment in gene expression involved in tissue reorganization and metabolic processes (Figure 1H) indicating that *Prkar1a* gene inactivation resulted in complex and deep disruption of adult testis functions and somatic cell tumor formation.

**Figure 1.**
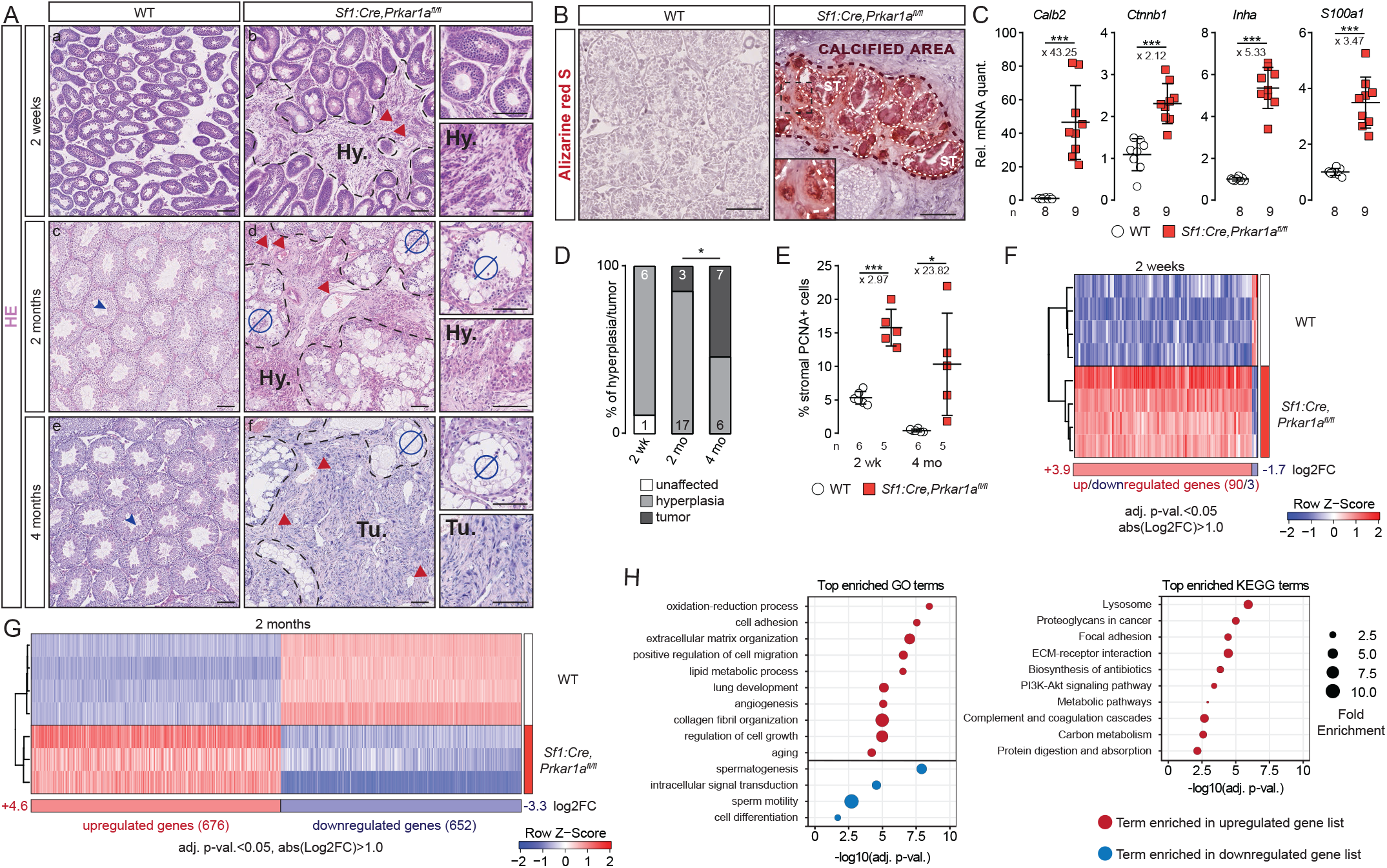
*Prkar1a* loss in mouse testicular somatic cells induces LCCSCT-like lesions. (**A**), HE staining of 2-wk (a, b), 2-mo (c, d) and 4-mo-old (e, f) WT and *Prkar1a* somatic progenitor cell knockout testes (noted as prKO, *Sf1:Cre,Prkar1a^fl/fl^*). Red arrowheads indicate prKO hyper-vascularization. Hy., hyperplasia; Tu., tumor; Ø, absence of spermatozoa. (**B**), Alizarine Red S staining in 2-mo-old prKO testis. Dashed lines delineate the calcium accumulation areas both in ST and stroma (inset). (**C**), RTqPCR analysis for LCCSCT markers transcripts (*Calb2*, *Ctnnb1*, *Inha*, *S100a1*) in 4-mo-old WT and prKO testes. Statistical analysis was performed using Student’s t-test or Welch’s t-test. (**D**), Relative proportion of hyperplasia and tumor from 2-wk-old to 4-mo-old prKO testes evaluated following HE staining. Number of each histological defect is indicated in bars. Statistical analysis was performed using two proportion Fisher’s test (hyperplasia/tumor proportions between 2-mo and 4-mo-old prKO testes). *p=0.0259*. (**E**), Stromal PCNA+ cells quantification represented as a percentage of cells in 2-wk and 4-mo-old WT and prKO testes. Statistical analysis was performed using Welch’s t-test. (**F**and **G**), Heatmap representing the median centered expression of significantly deregulated genes (abs(log2FC>1) % adj.p-val<0.05) in 2-wk (F) and 2-mo-old (G) prKO testes (n=3-4) compared with WT (n=4). FC, fold change. (**H**), Top 10 enriched GO and KEGG terms using 2-mo-old prKO significant deregulated genes. Bars represent the mean per group ± SD. Scale bars, 100μm. ns, not significant, *\*p<0.05*, *\*\*\*p<0.001.*

### Seminiferous tubules defects are associated with *Prkar1a* loss in Sertoli cells

*Sf1:Cre* mediated recombination in testis targeted both supporting and stromal cells and resulted in a complex phenotype in adult prKO mice with intratubular and stromal compartment alterations that could be amplified by reciprocal dysregulated paracrine signals. To unravel the cell specific *Prkar1a* loss impacts in the global LCCSCT pathogenesis, we generated lrKO (*Scc-Cre,Prkar1a^fl/fl^*) and srKO (Amh*-Cre,Prkar1a^fl/fl^*) mouse models with either Leydig or Sertoli cell-specific *Prkar1a* invalidation triggered from E13.5 and E15 respectively using *Cyp11a1:Cre* or *Amh:Cre* drivers (22, 23). The cell specificity of recombination was confirmed by analysing *Rosa26R-mtmg* reporter activity (Figure S4 A-D). Similarly to WT, 2-month-old lrKO mice were fertile and showed healthy testicular histology (Figure 2 A-E) devoid of stromal hyperplasia or tumor expansion. However like prKO mice, adult lrKO male mice developed hypertestosteronemia (Figure S4G). Transcriptomic data from comparative lrKO *vs* WT testes microarrays analyses confirmed fewer molecular deregulations in 2-month-old lrKO testes with only 22 significantly deregulated genes (17 up- and 5 down-regulated genes) (Figure 2G). By contrast loss of *Prkar1a* in SC (srKO model) was sufficient to reproduce intratubular and peritubular LCCSCT lesions and led to infertility (Figure 2 A-D) (Figure S4 E and F).The unbiased transcriptomic analyses revealed a huge deregulation of the molecular networks in 2-month-old srKO testes (560 deregulated, 150 up- and 410 down-regulated genes) (Figure 2F and G). Comparative heatmap analyses between WT, prKO, srKO, and lrKO model data highlighted a main cluster of 473 genes (84.4 % of srKO deregulated genes) exhibiting deregulations both in 2-month-old prKO and srKO mutant testes (114 up- and 359 down-regulated genes) compared with WT. Consistent with the germ cell population decrease, 75.5% of this cluster genes were significantly down-regulated and strongly associated with spermatogenesis alteration in gene set enrichment analyses (Figure 2G and H) (Figure 3A). Unlike prKO model, adult srKO mice had normal testosterone plasmatic levels and never developed tumour mass (up to 12 months of age) (Figure S4 G and H). In agreement with the delayed *Prkar1a* loss impact previously observed in prKO model, transcriptomic analyses of both 2-week-old lrKO and srKO testes revealed very few divergence from WT molecular signatures (Figure 2F). Altogether molecular and histological data from srKO mice demonstrated that loss of *Prkar1a* restricted to SC failed to initiate tumorigenesis although ST functions were altered.

**Figure 2.**
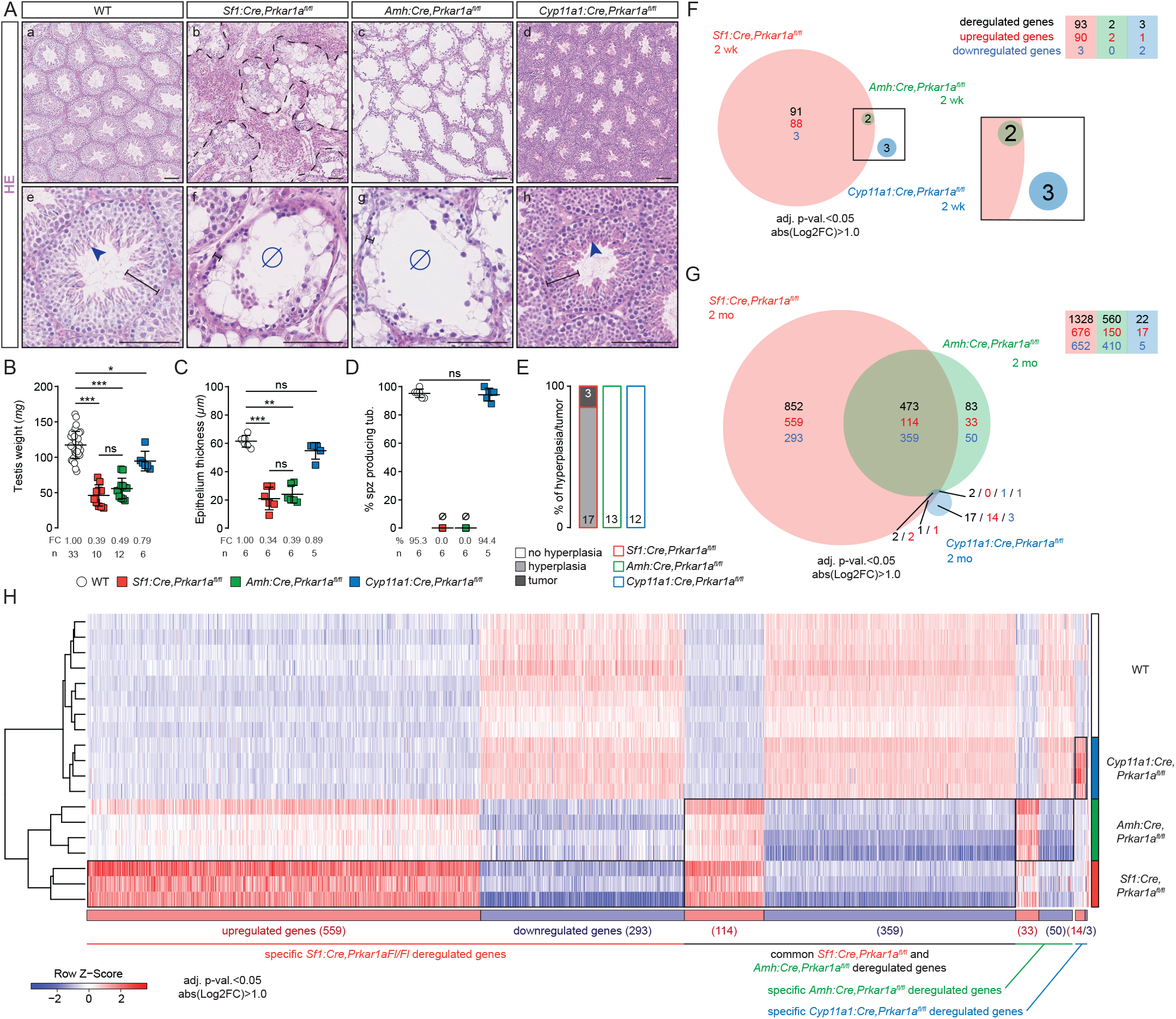
Seminiferous defects are associated with *Prkar1a* loss in Sertoli cells. (**A**), HE staining of 2-mo-old WT (a,e), prKO (*Sf1:Cre,Prkar1a^fl/fl^*; somatic progenitor cell *Prkar1a* inactivation) ((b,f), srKO (*Amh:Cre,Prkar1a^fl/fl^*; Sertoli cell *Prkar1a* inactivation) (c,g), lrKO (*Cyp11a1:Cre,Prkar1a^fl/fl^*; Leydig cell *Prkar1a* inactivation) testes (d,h). Dashed black lines delineate the tumor developed in prKO testis, blue arrowheads show spermatozoa and black lines limit seminiferous epithelium thickness. Ø, absence of spermatozoa. Scale bars, 100μm. (**B**), Testis weight of 4-mo-old WT, prKO, srKO and lrKO mice. One-way ANOVA was followed by Tukey multiple correction test. (**C**), Epithelium thickness quantified following HE staining in 2-mo-old WT, prKO, srKO and lrKO testes. Kruskal-Wallis’ test was followed by Dunn’s multiple correction test. (**D**), Quantification of ST with elongated spermatids represented as a percentage of positive tubules quantified following HE staining in 2-mo-old WT, prKO, srKO and lrKO testes. Statistical analysis was performed using Student’s t-test. (**E**), Relative proportion of hyperplasia and tumor in 2-mo-old prKO, srKO and lrKO testes. Numbers for each histological defect and genotype are indicated in bars. (**F** and **G**), Venn diagram of the significantly deregulated genes (abs(log2FC>1) % adj.p-val<0.05) in 2-wk-old (F) and 2-mo-old (G) prKO, srKO and lrKO testes compared with WT (n=3-4 mice per group). (**H**), Heatmap representing the median centered expression of significantly deregulated genes in 2-mo-old prKO, srKO and lrKO testes compared with WT. Bars represent the mean per group ± SD. *\*p<0.05*, *\*\*p<0.01*, *\*\*\*p<0.001.*

**Figure 3.**
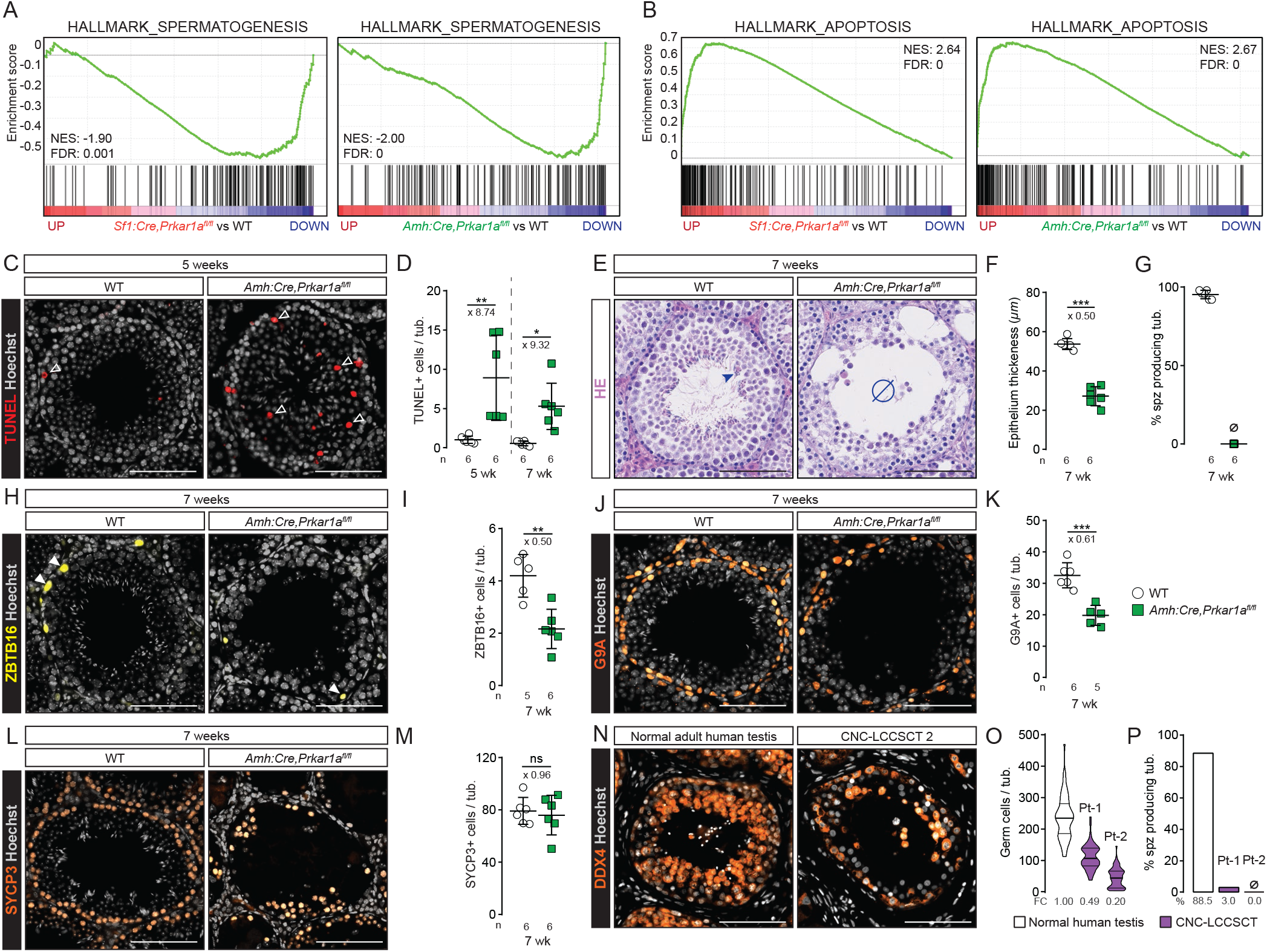
PKA overactivation in Sertoli cells triggers increased germ cell apoptosis from prepubertal age. (**A** and **B**), GSEA of microarray gene expression data showing a negative enrichment of hallmark-spermatogenesis gene list (A) and a positive enrichment of hallmark-apoptosis gene list (B) both in 2-mo-old prKO and srKO testes compared with WT. (**C**), TUNEL staining highlighting apoptotic cells in 5-wk-old WT and srKO testes. White arrowheads indicate TUNEL+ cells. (**D**), Quantification of TUNEL+ cells in 5-wk and 7-wk-old srKO testes compared with WT. Statistical analysis was performed using Welch’s t-test or Mann-Whitney’s test. (**E**), HE staining of 7-wk-old WT and srKO testes. Blue arrowheads, spermatozoa; Ø, absence of spermatozoa. (**F** and **G**), Epithelium thickness quantification (F) and percentage of ST with elongated spermatids (G) in 7-wk-old WT and srKO testes. Statistical analysis was performed using Student’s test. (**H**-**M**), Immunohistochemical detection and quantification of ZBTB16 (spermotogonia-A) (H and I), G9A (spermotogonia-B) (J and K) and SYCP3 (spermotocytes) (L and M) in 7-wk-old WT and srKO testes. White arrowheads indicate ZBTB16+ cells. Quantification is based on the number of positive cells per tube. Statistical analysis was performed using Student’s test. (**N**and **O**), DDX4 immunohistochemical detection (N) and germ cell loss quantification in CNC-LCCSCT compared with normal adult human testis (O). (**P**), Percentage of ST with elongated spermatids quantified following HE staining dropped in CNC-LCCSCT compared with normal adult human testis. Bars represent the mean per group ± SD. Scale bars, 100μm. *\*p<0.05*, *\*\*p<0.01*, *\*\*\*p<0.001.*

### PKA overactivation in Sertoli cell triggers increased germ cell apoptosis from prepubertal age

Adult prKO and srKO mice shared common deregulated molecular signatures associated with non-autonomous GC loss that could be attributed to disturbed Sertoli cells nursing capacities. Hallmark gene set enrichment analyses (GSEA) of a “spermatogenesis” gene set showed a robust negative enrichment in srKO and prKO testes, that was associated with an “apoptosis” gene set positive enrichment compared with WT (Figure 3, A and B). In order to specify the nature of the cellular events leading to the GC loss at adulthood in response of Sertoli *Prkar1a* inactivation, the developing GC types in srKO testes were evaluated. Significant increased TUNEL staining over the entire width of the germinal epithelium from 5-week and 2-month-old srKO testes confirmed that apoptosis induced by *Prkar1a* loss occurred at different spermatogenesis stages and contributed to decrease the germinal epithelium thickness (Figure 3, C-F). Although a complete lack of spermatozoa was evidenced in 2-month-old srKO ST (Figure 3G), immunostaining analyses inventoried only a significant 1.8-fold decrease in ZBTB16+ spermatogonia and a 1.6-fold decrease in G9A+ cells number (Figure 3, H-K) corresponding to the undifferentiated spermatogonia up to early leptotene spermatocyte populations. Moreover, SYCP3+ spermatocyte number was unchanged in srKO compared with WT (Figure 3, L and M), suggesting the occurrence of compensatory events counteracting increased apoptotic germ cell counts. Indeed, further time course analysis revealed a transient growing number of ZBTB16+ cells, G9A+ cells and SYCP3+ meiotic cells from 2 up to 5 weeks of age which paralleled a significant increased srKO testis weight from PND9 to 5 weeks compared with WT (Figure S5, A and B). This enhanced germ line expansion including elongated spermatid formation resulted in an abnormal full filling of the srKO ST as attested by testis weight that culminated at 5 weeks of age before being drastically drained thereafter (Figure S5, C-E). SOX9 immunostaining analyses revealed that SC number was significantly 1.17-fold increased in 2-week-old srKO testis compared with WT and 0.85-fold decreased at 7 weeks (Figure S5, F and G). However, these slight variations were unlikely to fully account for the magnitude of the alterations affecting the germ line. Altogether these data demonstrated that the lack of germ line in adult srKO testis was rather due to chronic apoptosis events overlapping deregulated increase of GC population than to a meiosis commitment blockade in response to SC *Prkar1a* gene inactivation. These observations are consistent with DDX4 immunostaining in human CNC-LCCSCT biopsies evidencing a strong decrease in GC number (Figure 3, N-P).

### PKA overactivation alters Sertoli cell polarity

The formation of flaking vacuoles inside the germinal epithelium associated with round clusters of GC released in the lumen was another phenomena specifically occurring in the testes of CNC patients and prKO and srKO mice that should result, together with apoptosis, in the loss of spermatogenic cells (Figure4A-D). Comparative time course analyses of srKO and prKO histological intratubular alterations indicated a more precocious prKO increase in vacuolated tubules into the germinal epithelium compared with srKO model that could contribute to worsen germ cell depletion (Figure 4D) (Figure S6A). Consistent with these observations, GSEA of the transcriptome of 2-month-old srKO testis revealed extended modification of the molecular networks supporting tubule architecture including, cytoskeleton, adhesion, or junctional proteins (Figure 4E). To gain insight into the modulators of the germinal epithelium integrity, we extracted expression levels of a list of 32 genes involved in Sertoli-Sertoli junctions or Sertoli-GC interactions. Fifteen of these 32 genes were significantly up-regulated and 3 genes were down-regulated both in srKO and prKO testes compared with WT (Figure 4F) and RT– qPCR conducted on independent WT, srKO and prKO samples confirmed their significative deregulated expression (Figure S6B). Immunohistological analyses of Claudin11 (*Cldn11*) and β-catenin (*Ctnnb1*) specified that these junction proteins involved in BTB establishment were delocalized to the entire membrane perimeter in both srKO and prKO mutant testes (5 wk) and that β-catenin staining was also diffused into the cytoplasm (Figure 4G-I). These delocalizations were associated with a SC cytoskeleton alteration attested by Tubulinβ3 densification and Vimentin disorganization (Figure S6C). The junctional defects were also present in the human CNC ST located out of the tumor mass as attested by Claudin11 and β-catenin IHC analyses (Figure 4J-L). These data suggested a compromised setting of the adluminal compartment by the blood-testis barrier (BTB) junctions which is critical to spermatogenesis achievement (24). Therefore BTB functional integrity has been tested in the srKO and prKO mutant mice by intratesticular injection of a fluorescent tracker (Figure 4, M and N). The tracker diffused inside ST and confirmed a drastic loss of BTB sealing in both srKO and prKO (6 weeks of age) demonstrating that SC failed to provide an appropriate structural dynamic for the GC maturation across the seminiferous epithelium. Moreover, transcriptomic and RTqPCR analyses (srKO/prKO testes) highlighted an earlier and huge up-regulation of *Dpt* expression in 2-week-old prKO testes which could amplify the BTB alterations, the vacuole formation and the germ cell loss as it was demonstrated in transgenic mice overexpressing *Dpt* in SC (25) (Figure 4O).

**Figure 4.**
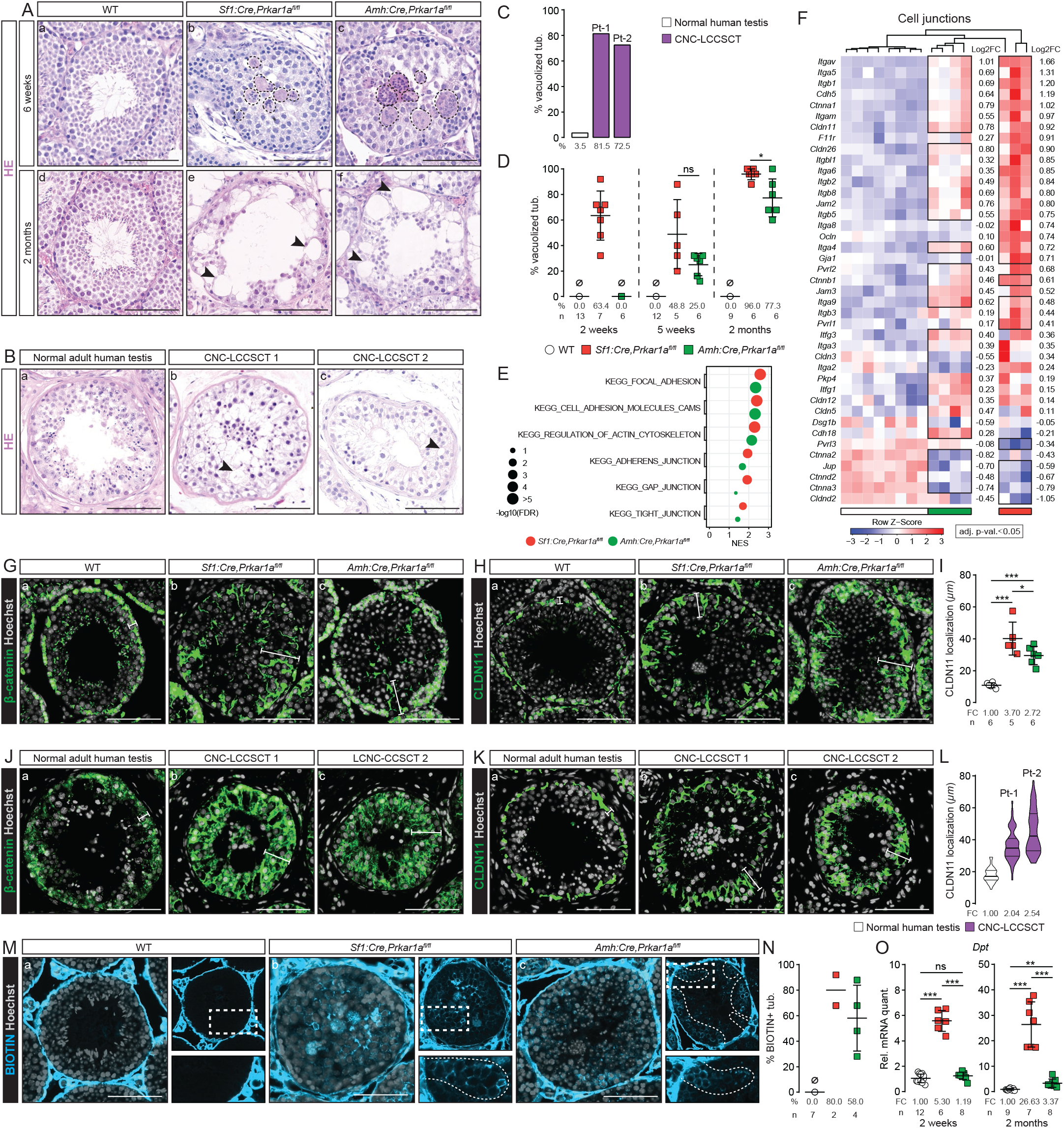
PKA overactivation alters Sertoli cell polarity. (**A**), HE staining of 6-wk (a-c) and 2-mo-old (d-f) WT, prKO and srKO testes. Black dashed lines delineate multinucleated giant cells at 6-wk resulting in vacuole formation (black arrowhead) at 2-mo. (**B**), Normal adult human testis (a) and CNC-LCCSCT (b,c) HE staining. (**C** and **D**), Quantification of vacuolized tubules represented as a percentage of positive tubules quantified following HE staining of normal adult human testis, LCCSCT CNC (C), 2-wk-old, 5-wk-old and 2-mo-old WT, prKO and srKO testes (D). Statistical analysis was performed using Welch’s t-test. (**E**), GSEA enrichment scores of microarray gene expression data using KEGG gene lists associated with junction/cytoskeleton in 2-mo-old prKO or srKO testes compared with WT. (**F**), Heatmap representing the median centered expression of significantly deregulated regulators of cell junctions (adj.p-val.<0.05) in 2-mo-old prKO and srKO testes (n=3-4) compared with WT (n=4). (**G**-**L**), Immunohistochemical detection of β-catenin (G and J), CLDN11 (H and K) and quantification of CLDN11 expression domain (I and L) based on the distance between basal lamina and apical CLDN11 accumulation in 5-wk-old WT, prKO, srKO testes (G-I) and in normal adult human testis and CNC-LCCSCT (J-L). White line delineates β-catenin or CLDN11 expression domain. One-way ANOVA was followed by Tukey multiple correction test. (**M**), Biotin tracer detection after injection underneath the testis capsule in 6-wk-old WT, prKO and srKO testes. White dashed line marks intra-seminiferous biotin diffusion in prKO and srKO testes. (**N**), Blood-testis barrier integrity loss evaluation based on the percentage of tube with intra-seminiferous biotin accumulation. (**O**), RTqPCR analysis of *Dpt* transcripts in 2-wk and 2-mo-old WT, srKO and prKO testes. Welch’s one-way ANOVA was followed by Games-Howell multiple correction test. Bars represent the mean per group ± SD.

### PKA overactivation modifies Sertoli signals involved in the control of spermatogenesis

The maintenance of GC within spermatogenic cycle is supported by mature SC through appropriate production of mitogens and differentiation factors that could be affected by *Prkar1a* gene inactivation. The interaction of the growth factor Kit-ligand (KIT-L) produced by SC with its specific c-KIT receptor expressed on GC from differentiating spermatogonia causes the PI3K-AKT pathway activation that is required for proliferation and survival of mitotic GC, both in the embryonic and postnatal testes (26–28). KIT-L also triggers PI3K/AKT and Ras/MAPK crosstalk that plays a critical role in the development of germ cell cancers (29). In addition, the KIT-L/c-KIT/JAK/STAT pathway is the main promoter of mast cell growth and differentiation from bone marrow and peripheral progenitors (30). Cyclic AMP/PKA-driven mechanisms leading to increased *Kitl* transcription in SC have been previously described (31, 32). Consistent with these previous observations, transcriptomic and RTqPCR analyses in prKO/srKO testes showed that *Prkar1a* loss led to a significant and early up-regulation of *Kitl* transcripts associated with a positive enrichment of a “reactome signaling by SCF (*alias* KIT-L)/KIT” gene set and to an increase in mast cells recruitment in interstitial tissue (Figure 5, A-D).

**Figure 5.**
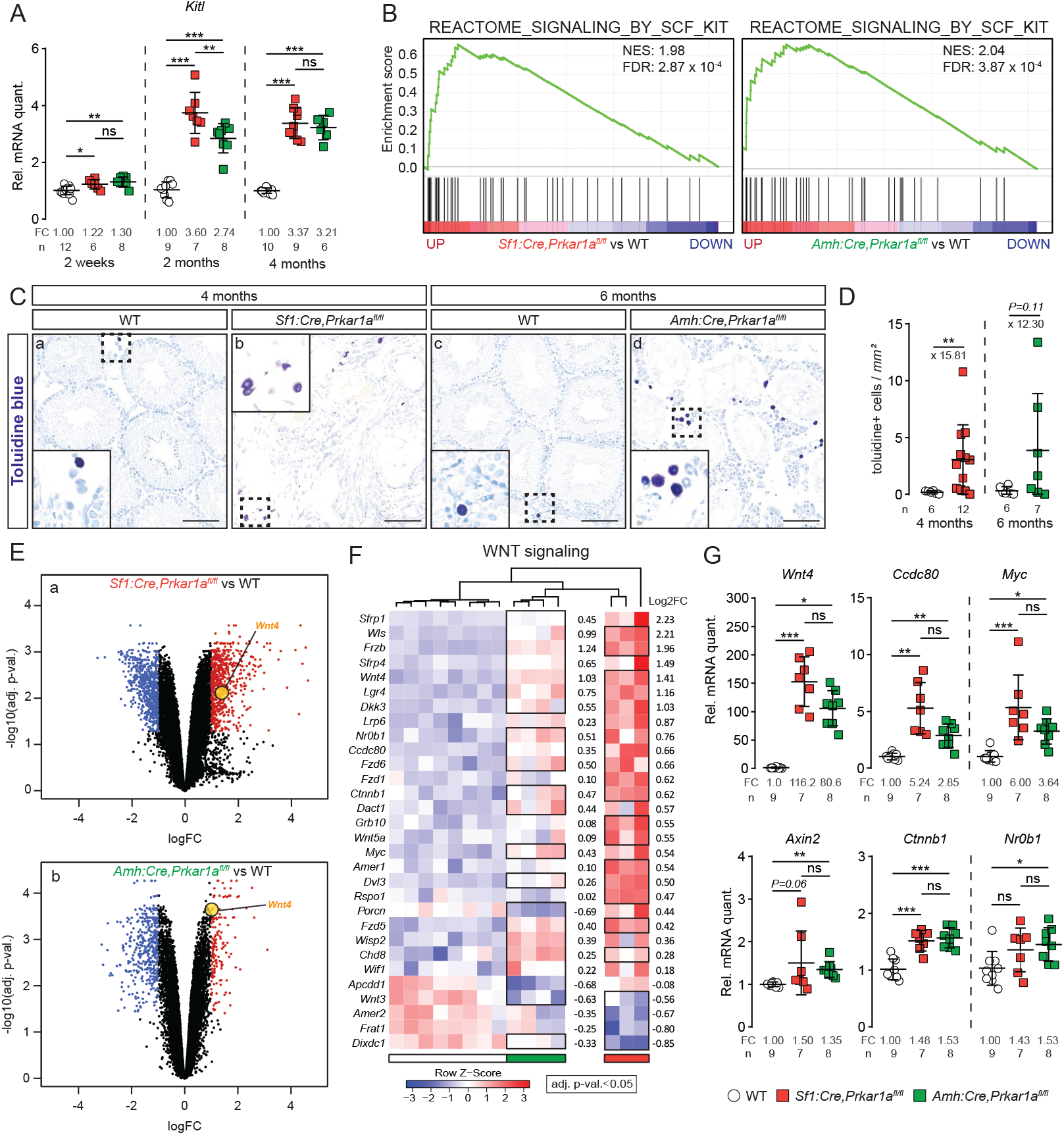
PKA overactivation modifies Sertoli signals involved in control of spermatogenesis. (**A**), *Kitl* transcripts RTqPCR analysis in 2-wk, 2-mo and 4-mo-old WT, prKO and srKO testes. One-way ANOVA was followed by Tukey multiple correction test and Welch’s one-way ANOVA was followed by Games-Howell multiple correction test. (**B**), GSEA of microarray gene expression data using REACTOME-signaling by SCF KIT gene list in 2-mo-old prKO and srKO testes compared with WT. (**C**), Toluidine blue staining showing mast cell recruitment in 4-mo-old prKO (a,b) and 6-mo-old srKO (c,d) testes. Scale bars, 100μm. (**D**), Mast cells quantification based on the toluidine+ cell number per histological section. Statistical analysis was performed using Welch’s t-test. (**E**), Volcano plot showing differential gene expression in 2-mo-old prKO (a) and srKO (b) testes compared with WT. Significantly deregulated genes (abs(log2F-C>1) % adj.p-val<0.05) are highlighted in blue (down) or in red (up). *Wnt4* (yellow point) up-regulated gene. (**F**), Heat-map representing the median centered expression of significantly deregulated WNT pathway regulators and target genes (adj.p-val.<0.05) in 2-mo-old prKO and srKO testes (n=3-4) compared with WT (n=4). (**G**), RTqPCR analysis of WNT pathway regulators (*Wnt4*, *Ctnnb1*) and target genes (*Axin2*, *Ccdc80*, *Myc*, *Nr0b1*) expression in 2-mo-old WT, prKO and srKO testes. One-way ANOVA was followed by Tukey multiple correction test, Welch’s one-way ANOVA was followed by Games-Howell multiple correction test and Kruskal-Wallis’ test was followed by Dunn’s multiple correction test. Bars represent the mean per group ± SD. *\*p<0.05*, *\*\*p<0.01*, *\*\*\*p<0.001.*

Surprisingly, further transcriptomic and RTqPCR analyses highlighted a significant up-regulation of the *Wnt4* gene expression that was listed in the common deregulated gene cluster from adult prKO and srKO models indicating that most of the WNT4 overproduction originated from adult SC in response to *Prkar1a* inactivation (Figure 5, E and F). Heatmap of a “WNT signaling” gene set showed a robust positive enrichment of these molecular signatures in adult srKO and prKO testes compared with WT, suggesting a functional significance of the WNT pathway activation. This observation was confirmed by RTqPCR analysis of *Axin2, c-Myc, Ccdc80* and *Nr0b1* target gene expression (Figure 5G) (33, 34). Altogether these data revealed that PKA constitutive activation leads to a new *Wnt4* up-regulation in Sertoli cells whereas PKA dependent *Wnt4* gene expression was only previously evidenced in other tissue differentiations such as kidney tubulogenesis or uterine stromal cells decidualization (35, 36).

### *Wnt4* inactivation decreases germ cell apoptosis induced by *Prkar1a* loss

WNT4/β-catenin signaling pathway was involved in stem cell proliferation of several tissues (37). Although WNT4/β-catenin signaling was shown to be essential modulator of female ovary differentiation (38), *Ctnnb1* expression was dispensable for testis development (39). Moreover, Boyer A, Yeh JR et al, (2012) demonstrated that excessive WNT4 signal repressed the normal activity of germ stem cells by inducing their apoptosis (40), while forced expression of active β-catenin in fetal gonocytes deregulated spermatogonial proliferation and apoptosis (41, 42). Our analyses indicated that the increase GC apoptosis together with disturbed BTB establishment impaired the spermatozoa accumulation in prKO and srKO mice. In order to assess the role of WNT4 in GC altered homeostasis triggered by *Prkar1a* gene loss, we generated double KO mice for *Prkar1a* and *Wnt4* in somatic cells (prwKO model; *Sf1:Cre,Prkar1a^fl/fl^,Wnt4^fl/fl^*) or restricted to Sertoli cells (srwKO model; *Amh:Cre,Prkar1a^fl/fl^, Wnt4^fl/fl^*) (Figure S7, A and B). As expected, simple mutant mice bearing targeted *Wnt4* inactivation in SC (swKO) or in somatic cells (pwKO) were fertile and had a normal testicular development and physiology (Figure S7C). Unlike the srKO testis, ST from 8-week-old double mutant srwKO testis maintained a complete germinal epithelium containing all GC types including spermatozoa, associated with a significant decrease in intratubular apoptosis events compared with srKO testes (Figure 6, A-J). This indicates that *Wnt4* loss in a PKA activation context allowed the achievement of a complete GC lineage whereas SC junction disorganization, flacking vacuole and infertility were still maintained (Figure 6, J-M). Therefore, SC-specific PKA constitutive activity up-regulated the *Wnt4* expression which increased germ cell apoptosis. Nevertheless, several spermatocytes with abnormal mitotic features not only persisted in srwKO but also were accentuated in prwKO testis despite normalization of the germ line marker gene *Ddx4 (*Figure 6A, insets) (Figure S7D). These data revealed a critical role of SC PKA pathway on spermatogenesis independent from WNT4 signal that impaired long-term maintenance of GC. Transcriptomic and RTqPCR analyses highlighted increased *Ctnnb1* mRNA levels in the adult testes of both prKO and srKO mice that remained elevated in srwKO and prwKO (Figure S7E) demonstrating that PKA induced *Ctnnb1* expression independently from WNT4 signal.

**Figure 6.**
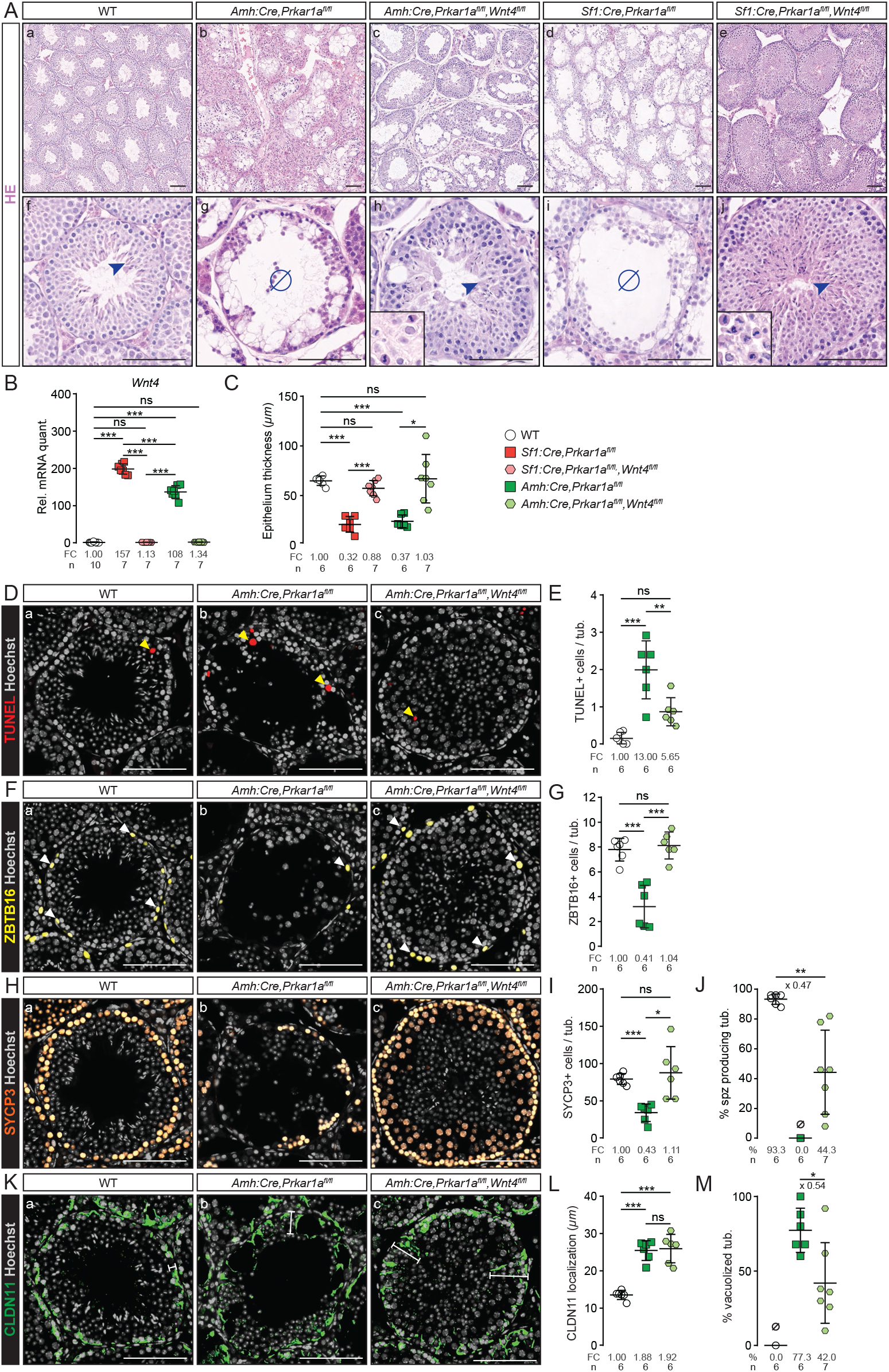
*Wnt4* inactivation decreases germ cell apoptosis induced by *Prkar1a* loss. (**A**), HE staining of 2-mo-old WT (a,f), prKO (b,g), double mutant prwKO testes (*Sf1:Cre,Prkar1a^fl/fl^,Wnt4fl/fl*) (c,h), srKO (d,i) and srwKO testes (*Amh:Cre,Prkar1a^fl/fl^,Wnt4^fl/fl^*) (e,j). Arrowheads, spermatozoa; Ø, absence of spermatozoa; insets, abnormal mitotic features. (**B**), RTqPCR analysis of *Wnt4* transcripts in 2-mo-old WT, prKO, prwKO, srKO and srwKO testes. Welch’s one-way ANOVA was followed by Games-Howell multiple correction test. (**C**), Epithelium thickness quantified following HE staining in 2-mo-old WT, prKO, prwKO, srKO and srwKO testes. Welch’s one-way ANOVA was followed by Games-Howell multiple correction test. (**D**-**E**), TUNEL staining in 2-mo-old WT, srKO and srwKO testes (D) and TUNEL+ cells quantification (E) in 2-mo-old srwKO testis compared with WT and srKO testes. Yellow arrowheads indicate TUNEL+ cells. One-way ANOVA was followed by Tukey multiple correction test. (**F**-**I**), Immunohistochemical detection of ZBTB16 (F) and SYCP3 (H) in 2-mo-old WT, srKO and srwKO testes and quantification of ZBTB16 (G) and SYCP3 (I) based on the number of positive cells per tube. White arrowheads indicate ZBTB16+ cells. Welch’s one-way ANOVA was followed by Games-Howell multiple correction test. (**J**), Percentage of ST with elongated spermatids quantified following HE staining in 2-mo-old WT, srKO and srwKO testes. Statistical analysis was performed Welch’s t-test. (**K**), Immunohistochemical detection of CLDN11 in 2-mo-old WT, srKO and srwKO testes. (**L** and **M**), Quantification of CLDN11 expression domain (L) and percentage of vacuolized tubules (**M**) in 2-mo-old srwKO testis compared with WT and srKO. Statistical analysis was performed using Student’s t-test or one-way ANOVA followed by Tukey multiple correction test. Bars represent the mean per group ± SD. Scale bars, 100μm. *p<0.05, **p<0.01, ***p<0.001.

### *Wnt4* inactivation impaired tumor mass formation induced by *Prkar1a* loss

The human LCCSCT (CNC) tumor masses are the site of proliferative activity mediated by misunderstood mechanisms. Consistent with LCCSCT in CNC context, from 8 weeks of age, prKO testis developed early proliferative tumour masses arising from neonatal stromal hyperplasia. RTqPCR analyses highlighted an up-regulation of *Wnt4* and *Ctnnb1* expression in 2-month-old prKO mice suggesting, together with WNT signaling heatmap analysis (Figure 5, F and G) that a PKA induced WNT/β-catenin pathway could be involved in the LCCSCT promotion in CNC patients and prKO mice. In addition to the *Wnt4* up-regulation, further analyses indicated that the loss of *Prkar1a* led to a chronic increase of both active β-catenin accumulation and gene expression of *Fzd6* and *Wnt5a* in 4-month-old prKO testes (Figure 5F) (Figure 7, A-C) that were previously demonstrated to exacerbate tumor expansion through canonical and non-canonical WNT pathways activity in various cancers (43). Histological examinations of mutant testes revealed that although the oldest double mutant prwKO testes were still the site of stromal fibrosis, they were devoid of tumor mass compared with single mutant prKO testes (Figure 7, D-F) (Figure S7E). Comparative PCNA IHC analyses confirmed that *Wnt4* inactivation strongly decreased the stromal proliferation that was triggered by *Prkar1a* loss in somatic testis cells (Figure 7, G and H). Moreover, whole testis RTqPCR analyses confirmed that in these prwKO testes, *Ccdc80*, *Axin2* and *Ki67* transcripts were reduced compared with 4-month-old prKO testes (Figure 7, I and J). All together these results demonstrate that WNT4 is a testis tumor signal in response to *Prkar1a* inactivation. Consistent with our murine models, IHC analyses confirmed that human CNC-LCCSCT tumor mass cells retained WNT4 huge staining that colocalized with cytoplasmic β-catenin accumulation. This demonstrates that WNT/β-catenin activation is likely associated with the human CNC-LCCSCT expansion (Figure 7K).

**Figure 7.**
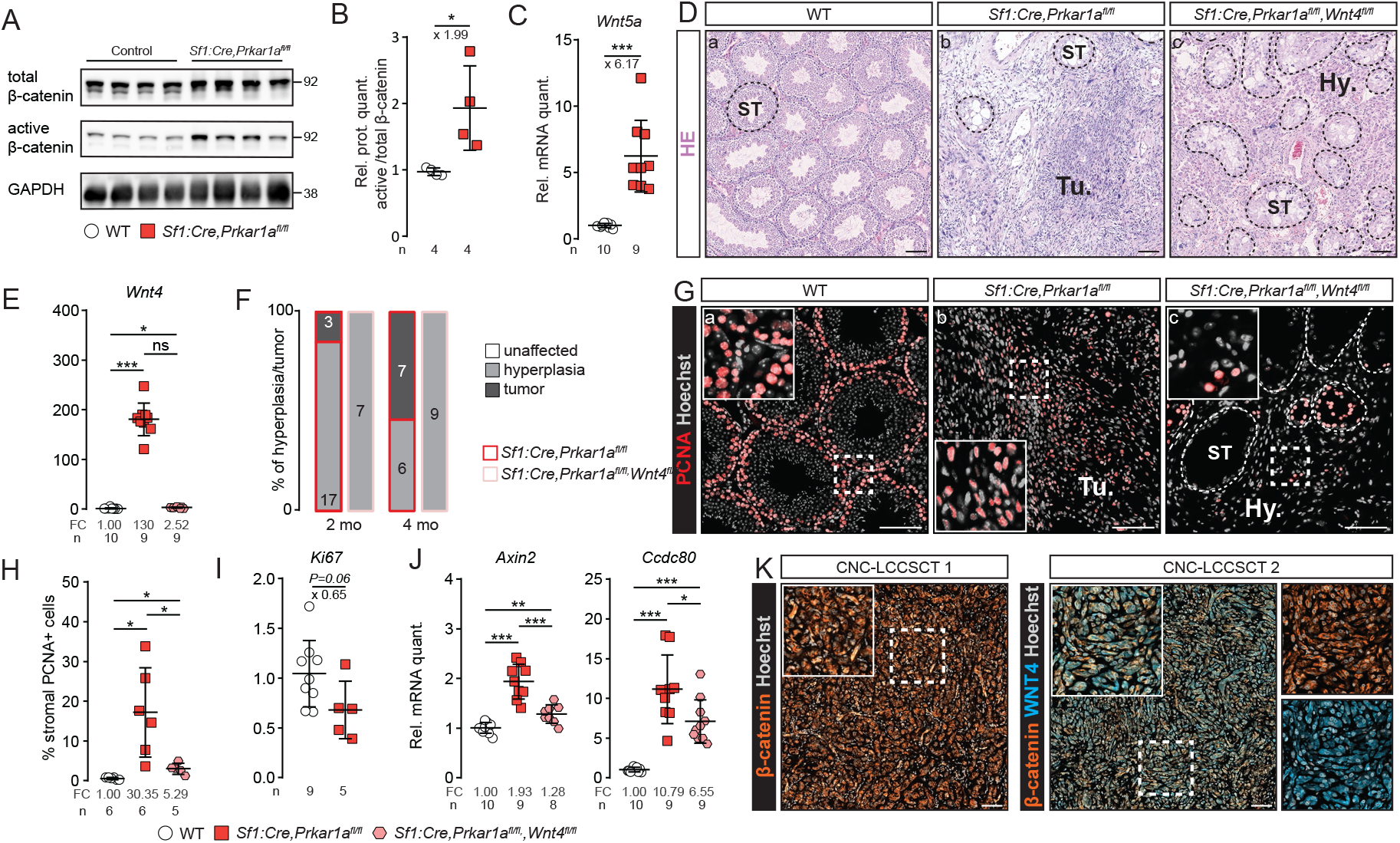
*Wnt4* inactivation impairs tumor mass formation induced by *Prkar1a* loss. (**A**), Active β-catenin accumulation in 4-mo-old WT and prKO testes. (**B**), Active β-catenin quantification over total β-catenin. Statistical analysis was performed using Student’s t-test. (**C**), *Wnt*5a transcripts in 4-mo-old WT and prKO testes. Statistical analysis was performed using Welch’s t-test. (**D**), HE staining of 4-mo-old WT, prKO and prwKO testes. Dashed lines delineate ST. Tu., tumor; Hy.,hyperplasia. (**E**), *Wnt4* transcripts in 4-mo-old WT, prKO and prwKO testes. One-way ANOVA was followed by Tukey multiple correction test. (**F**), Relative proportion of testicular hyperplasia and tumor in 2-mo and 4-mo-old prKO and prwKO testes was evaluated following HE staining. Numbers for each histological defect and genotype are indicated in bars. (**G**), Immunohistochemical analysis of PCNA in 4-mo-old WT, prKO and prwKO testes. (**H**), Quantification of stromal PCNA+ cells represented as a percentage of cells in 4-mo-old WT, prKO and prwKO testes. Welch’s one-way ANOVA was followed by Games-Howell multiple correction test. (**I**), *Ki67* transcripts RTqPCR analysis in 4-mo-old prKO and prwKO testes. Statistical analysis was performed using Student’s t-test. (**J**), *Axin2* and *Ccdc80* transcripts RTqPCR analysis in 4-mo-old WT, prKO and prwKO testes. One-way ANOVA was followed by Tukey multiple correction test and Welch’s one-way ANOVA was followed by Games-Howell multiple correction test. (**K**), Coimmunostaining for β-catenin and WNT4 in CNC-LCCSCT. Bars represent the mean per group ± SD. Scale bars, 100μm. *\*p<0.05*, *\*\*p<0.01*, *\*\*\*p<0.001.*

### PKA constitutive activation in somatic cells activates growth factor signaling pathways in CNC-LCCSCT

Unbiased gene ontology and GSEA highlighted additional prKO specific enrichment in signatures associated with proliferative pathways that could participate in tumor formation including up-regulated expression for *Fgf2, Egf, Igf1,* and *Tgfb1-3* (Figure 8, A-D). RTqPCR confirmed *Fgf2* up-regulation in prKO testes compared with WT, that was reduced in prwKO suggesting a possible *Wnt4* dependent involvement of the FGF2 signaling pathway in tumour development. By contrast increased *Egf*, *Igf1* and *Tgfb1-3* transcript levels in prKO testes remained unaffected by *Wnt4* loss suggesting that their PKA induced up-regulation was not sufficient to trigger precocious tumor formation (Figure 8E). Proliferative/oncogenic activities of the growth factors (IGF1, EGF, FGF2, KIT-L) overexpressed in prKO testes are mainly mediated by converging signaling pathway including RAS/MAPK and PI3K/AKT/mTOR signaling pathways (44–47). Immunodetection of activated MAPK modulators and mTOR targets in WB revealed significant increased levels of phosphorylated forms of ERK, 4EBP1 and S6K in 4-month-old prKO testes compared with WT (Figure 8, F and G) confirming the prKO gene enrichment analyses. In testis, the mTOR pathway was previously shown to control SC polarity and spermatogenesis (48, 49). Additionally its overactivity was demonstrated to mediate SC disruption in mice model lacking *Lkb1* gene in Sertoli cells (50). In prKO testes, IHC analyses revealed increased p4EBP1 accumulation in tubular compartment but also in tumor mass compared with WT, demonstrating that mTOR pathway was activated in tumor cells in response of *Prkar1a* loss (Figure 8H). Consistent with these observations, IHC analyses revealed that the human LCCSCT samples retained strong p4EBP1 immunostaining restricted to the majority of tumor cells and absent from non-tumor areas (Figure 8I). Moreover, to a lesser extent the human LCCSCT masses also displayed numerous foci of pERK accumulation demonstrating that these two proliferative pathways are activated in CNC-LCCSCT in response to *PRKAR1A* inactivation (Figure 8J). LCCSCT are solid tumors, this implies they could be the site of hypoxia. This hypothesis was strongly suggested in 2-month-old prKO testes by GO and GSEA analyses highlighting positive enrichment in hypoxia responsive genes (Figure 8, A and K). RTqPCR analyses in 4-month-old prKO mice confirmed the increased mRNA levels for *Hif1a, Hif1b,* and *Hif2a* genes that played a central role in O2-dependent transcriptional response (Figure 8L). Consistent with these observations, IHC analyses in human LCCSCT revealed increased cytoplasmic and nuclear HIF2A protein accumulation restricted to the tumor mass cells (Figure 8M). Altogether our data reveal that LCCSCT are the site of multiple proliferative pathways including mTOR, MAPK and hypoxia, acting with active WNT/β-catenin pathway to promote growth tumor.

**Figure 8.**
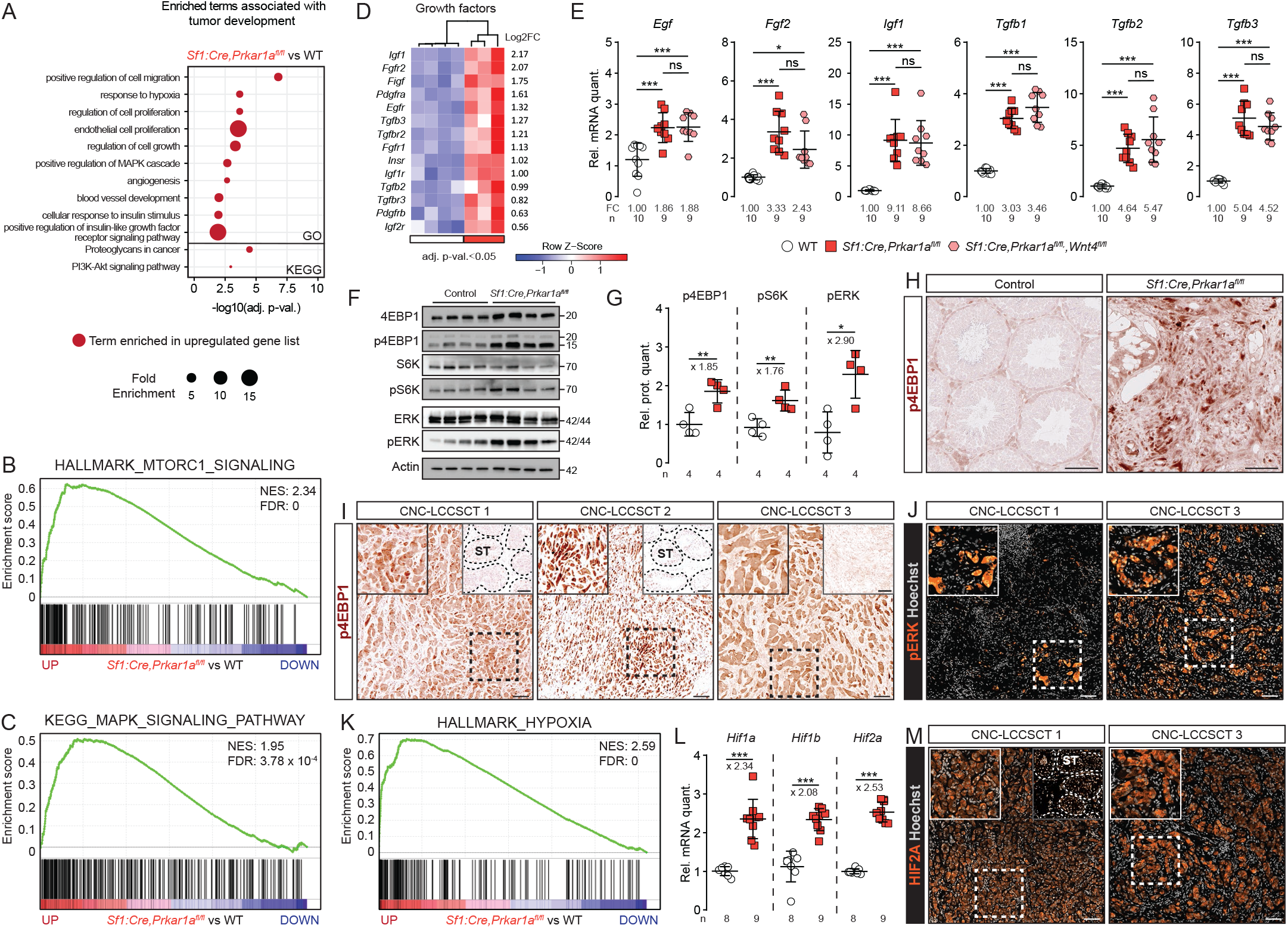
PKA activation induces mTOR, MAPK and hypoxia pathways in tumor of human LCCSCT. (**A**), Gene ontology enrichment scores of tumor development associated pathways based on gene list specifically deregulated in 2-mo-old prKO testis compared with srKO and lrKO testes. (**B** and **C**), GSEA of microarray gene expression data showing positive enrichment of hallmark-MTORC1 signaling gene list (B) and KEGG-MAPK signaling pathway gene list (C) in 2-mo-old prKO testis compared with WT. (**D**), Heatmap representing the median centered expression of significantly deregulated growth factors and associated receptors (adj.p.val.<0.05) in 2-mo-old prKO (n=3-4) compared with WT (n=4). (**E**), RTqPCR analysis of *Egf*, *Fgf2*, *Igf1*, *Tgfb1*, *Tgfb2* and *Tgfb3* transcripts in 4-mo-old WT, prKO and prwKO testes. One-way ANOVA was followed by Tukey multiple correction test, Welch’s one-way ANOVA was followed by Games-Howell multiple correction test and Kruskal-Wallis’ test was followed by Dunn’s multiple correction test. (**F**), Western blot analysis of p4EBP1, pS6K and pERK accumulation in 4-mo-old WT and prKO testes. (**G**), Phosphorylated protein quantification over total proteins. Statistical analysis was performed using Student’s t-test. (**H**), Immunohistochemical detection of p4EBP1 in 4-mo-old WT and prKO testes. (**I**and **J**), Immunohistochemical detection of p4EBP1 and pERK in CNC-LCCSCT. (**K**), GSEA of microarray gene expression data showing positive enrichment of hallmark-hypoxia gene list (genes upregulated in response to hypoxia) in 2-mo-old prKO testis compared with WT. (**L**), *Hif1a*, *Hif1b* and *Hif2a* transcripts analysis in 4-mo-old WT and prKO testes. Statistical analysis was performed using Welch’s or Student’s t-test. (**M**), Immunohistochemical detection of HIF2A in CNC-LCCSCT. Bars represent the mean per group ± SD. Scale bars, 100μm. *\*p<0.05*, *\*\*p<0.01*, *\*\*\*p<0.001.*

## DISCUSSION

To gain a better understanding of the CNC-LCCSCT pathogenesis, we performed *Prkar1a* inactivation either targeted in the entire somatic compartment or in main sub-types of testicular somatic cells in mice. These unique mouse models, offering a more homogeneous sampling than human biopsies, provide access to the early stages of the pathology as well as to advanced stages of tumor development. In addition, they constitute unvaluable tools for deciphering crosstalks between deregulated molecular circuits.

The prKO mice develop all the alterations identified in CNC-LCCSCT patients (bilateral tumors arranged in solid cords, fibrosis and calcifications associated with the increased expression of sex cord tumor markers (*Calb2, Ctnnb1, Inha and S100*)) and therefore they constitute a reliable model for exhaustive analyses of the molecular alterations caused by the lack of R1α subunit.

The ontogenic analysis of prKO model showed that major changes in endocrine activities, histological features and testicular transcriptome did not appear before the pre-pubertal stage, thus with a delay compared with early fetal inactivation of *Prkar1a* gene. This suggests that lack of R1α subunit has little impact on testicular ontogeny during the fetal and perinatal period. Conversely in the *princeps* CNC mouse model carrying constitutive homozygous *Prkar1a* deletion, lethality occurred as early as E8.5 due to growth defects in mesoderm-derived cardiac tissues which can be rescued by further inactivation of *Prkaca* gene encoding the Cα catalytic subunit of PKA (10). Possible compensatory mechanisms involving other PKA regulatory subunits, PKA inhibitor peptides or phosphodiesterases, in the developing fetal prKO testis remain to be determined. It should be noted that, although the occurrence of LCCSCT in CNC patients may be observed early in their first decade (5, 6, 51), no case of testicular abnormalities have been reported at birth, suggesting that in humans, as in our mouse models, the *Prkar1a* mutation does not cause abnormal fetal gonadal development. In our models, the total absence of tumor in the double mutant testis *Sf1:Cre,Prkar1a^fl/fl^,Prkaca^+/−^*, demonstrates that the induction of testicular lesions of prKO mice relies on the unrepressed catalytic activity of PKA rather than on a defect of the R1α subunit *per se*, in agreement with previous observations in other tissue lesions of the CNC (9, 52, 53).The prKO testes transcriptomic analysis revealed a modified expression profile for more than 1328 genes in the 2-month-old testis associated with a tissue and functional homeostasis disruption which reflects the complexity of the mechanisms resulting from the *Prkar1a* loss in all gonadal somatic cells. In addition, molecular signatures and histological analyzes revealed germline damages resulting from paracrine/non autonomous cell actions. Only very few studies have looked at the infertilities associated with *PRKAR1A* mutations in CNC patients, nevertheless they brought out alterations in male GC (53, 54) as consequences from *PRKAR1A* loss in somatic cells as well as in GC. The comparison of gene signatures of prKO testis with those of lrKO and srKO models allowed to demonstrate that *Prkar1a* inactivation restricted to Sertoli cells led to a progressive germ line loss from the prepubertal stages resulting in infertility at puberty. In both prKO and srKO ST, the underlying mechanisms include an increase in GC apoptosis and formation of GC syncytia containing flaking vacuoles reminiscent to those observed at the LCCSCT periphery in CNC patients. These massive apoptosis and desquamation mask the exacerbated increase in germ population that occurs in the first spermatogenesis waves in srKO testis compared with littermate control mice. These apoptotic events differ from the phenotypes observed in mouse models bearing FSHR signaling up-regulation (tgFSH/hpg and MT-hFSHR mice) in which at best, only an enhancement of spermatogonia proliferation and meiotic/post-meiotic cells survival were observed (55, 56). Likewise, the activating mutations of FSH signaling described in human species do not induce any testicular phenotype nor affect the fertility of male patients (57, 58). While it is well documented that FSH/FSHR/PKA pathway exerts its effects on the germ line *via* the production of SC growth signals such as FGF2, GDNF or KIT-L (31, 59–63, 32), our data indicate that constitutive activation of PKA in the SC triggers additional circuits that oppose actions normally arising from FSH/FSHR signaling. Indeed, the transcriptomic analysis of prKO and srKO testes allowed us to point out a very strong WNT4 signal overproduction by mutant SC. This observation is reminiscent to a previously described mouse model showing that SC-specific β-catenin constitutive activation leads to WNT4 overproduction and subsequent decrease in the spermatogonial stem cells activity responsible for apoptosis and loss of GC (40). Interestingly, in response to *Prkar1a* inactivation, our models also show WNT/β-catenin activation attested by up-regulation of *Wnt4* and *Ctnnb1* expression and by increased accumulation of the β-catenin active form (*i.e.* dephosphorylated). In CNC patients, we also observed intracytoplasmic accumulation of β-catenin in Sertoli cells from testicular biopsies. Moreover, we found that about 36% of the genes up-regulated in Sertoli cells from the *Ctnnb1^tm1Mmt/+^;Amhr2^tm3(cre)Bhr/+^* model (40) are also up-regulated in srKO testis indicating the establishment of partially common molecular alterations resulting from the mobilization of the WNT/β-catenin signaling pathway in the 2 models. Finally, *Wnt4* inactivation, suppressing germ line apoptosis in srwKO mice, definitely confirms the hypothesis of a converging phenotype between these different mutant models, and demonstrates the cooperation between overactive PKA and WNT/β-catenin signaling in triggering germ line alteration. However, unlike the observations from Boyer and coll., our analyzes of prKO and srKO mutants indicate that the PKA-induced *Wnt4* up-regulation is not associated with acquisition of granulosa *Fst* marker. Moreover SC maintain high *Sox9* expression while both *Foxl2* and *Rspo1* expression are absent from all *Prkar1a* mutant testis (40). Altogether, the comparison of *Ctnnb1^tm1Mmt/+^;Amhr2^tm3(cre)Bhr/+^* and prKO or srKO models suggests that constitutive activation of PKA both triggers massive overexpression of *Wnt4* and opposes its female identity programming function probably by reinforcing the maintenance of pro-testicular factors such as SOX9 which regulates its own expression and represses *Foxl2* expression (16, 64–67). Transcriptomic and IHC analyses reveal an increased production of the BTB proteins and SC associated cytoskeleton components in both mutant murine and CNC human ST. These junctional proteins are known to be positively controlled by various factors including SOX9, but also androgen and mTOR signaling pathways that are enhanced in our models (49, 64, 68). However, an alteration of GC progression into the adluminal tubular compartment due to paracellular extension of sertolian BTB proteins is unlikely to contribute to GC apoptosis in *PRKAR1A*/*Prkar1a* mutant human and murine testes. Indeed, in double mutant prwKO ST, the extended distribution of BTB proteins persists, while the GC apoptosis blockade partially restores the accumulation of elongated spermatids. This indicates that structural changes in the BTB induced by *Prkar1a* loss are mostly independent from WNT4 action and do not prevent spermatogenesis achievement. Nevertheless, the SC nuclei delocalization and the BTB functional failure are loss of polarity criteria suggestive of SC maturity disturbance in the srKO and prKO testes. However, unlike the *Ctnnb1^tm1Mmt/+^; Amhr2^tm3(cre)Bhr/+^* model (69), SC do not proliferate in the srKO and prKO ST and no overproduction of AMH persists at adulthood. These data indicate that activation of PKA in Sertoli cells induces specific pro-differentiating actions that counteract some exacerbated effects of WNT/β-catenin signaling observed in mouse models developed by Tanwar and coll. and Boyer and coll (40, 69). In particular, the *Prkar1a* mutant SC early display much higher increase of *Inha* transcripts than any other TGFβ family genes. This could play a major role in limiting the SC proliferation induced by the WNT/β-catenin pathway, as suggested by previous studies on inhibinα autocrine actions (70, 71). In addition, previously defined molecular signatures attesting the Sertoli cells maturation under the control of androgens (68, 72, 73) and TH (74, 75) remain unchanged in srKO and prKO transcriptomes at the pre-pubertal stage (2 weeks) but increase at adulthood. Finally, whether PKA-induced *Wnt4* expression results from PKA-dependent increased *Ctnnb1* expression and/or PKA-mediated β-catenin activation mechanisms as described in ovarian granulosa and luteal cells (76–78) remains to be explored for the *Prkar1a*^−/−^ SC. By contrast to the srKO and lrKO models, the prKO mice develop testicular tumors with similar histological and molecular characteristics to LCCSCT of CNC patients. This indicates that tumor onset is dependent on both SC and stromal proliferative signals activated by *Prkar1a* loss in both testicular compartments. This unprecedent situation in genetic models of testicular tumors underlines the role of paracrine interactions in the CNC-LCCSCT pathogenesis. Among the up-regulated signals in response to *Prkar1a* loss, *Wnt4* expression is present at significantly higher levels in the prKO than in the srKO model and increases in parallel with the tumor development suggesting that tumor cells could be an additional source of WNT4 signal. Indeed, our IHC analyses on human CNC-LCCSCT biopsies confirm that WNT4 is accumulated into the cells of the tumor mass. This increase in the WNT4 signal is associated with an increase in the expression of WNT/β-catenin responsive genes and therefore raised the question of its involvement in tumor promotion of LCCSCT. The double mutant prwKO model study demonstrates for the first time that the WNT4 signal is a limiting component in the growth factors set that conditions the CNC-LCCSCT development. Indeed, in the prwKO testis, the *Wnt4* loss abolishes the tumor formation and drastically reduces the stromal proliferation rate compared with prKO testis. Importantly, the *Wnt4* loss does not prevent the overall WNT/β-catenin pathway activation in the prwKO testis, given that the WNT5A ligand remains produced in excess, and that PKA-dependent phosphorylation can directly favor β-catenin stabilization (79). In this context, it can be assumed that *Wnt4* inactivation sufficiently reduces the strength of the pathway to limit stromal proliferation below a limiting threshold for tumor formation. Residual stromal hyperplasia in the double mutant testis also confirms the impact of additional PKA-activated growth signals independently from *Wnt4* expression. Previously, a mouse model (*Ctnnb1^tm1Mmt/+^;Tg(AMH-cre)1Flor*) expressing a stabilized form of β-catenin targeted in Sertoli cells developed testicular cancers with molecular and invasive characteristics different from tumors of the prKO model and human CNC-LCCSCT and which arise independently of the influence of stromal compartment (80). In human XY, while *WNT4* duplications have been associated with 46, XY testicular dysgenesis and ambiguous external genitalia occurring from fetal development, and despite the recurrent *CTNNB1* activating mutations in Sex Cord Stromal Tumor (SCST) cells (81), *WNT4* activating mutations remains poorly documented in testis cancer incidence unlike other tissues including ovaries (82). These observations reinforce the idea that although WNT4 signal is a mandatory actor, it does not support on its own the formation of the LCCSCT.

Comparative analyses of the prKO and srKO testis transcriptomes at 2 months suggested an overproduction of other proliferation signals in the prKO tumor among which FGF2 signal is of particular interest since *Fgf2* mRNA levels are closely associated with the tumor and decrease with *Wnt4* inactivation in prwKO testis. In healthy testis, *Fgf2* expression was shown to be responsive to FSH in SC (59) and to be expressed in Leydig cells where it promotes Leydig stem cell proliferation and commitment (83). Moreover, our analyzes indicate that the prKO testes exhibit increased expression of the *Fgfr1, Fgfr2*, and *Scd1* genes compared with srKO and WT testes suggesting the coregulation of these key partners in appropriate amounts to optimize transduction of the FGF2 signaling in the tumor (84). In agreement with a possible FGF2 involvement in human testis cancer, a study pointed significant increases in serum FGF2 levels in patients with testis tumors associated with positive expression inside tumors biopsies (85). Although, *Egf, Igf1 and Tgfb1-3* gene expressions were also strongly up-regulated in prKO testes, they remained elevated in prwKO testes, suggesting that they could participate in maintaining the moderate proliferation observed in the prwKO residual stromal hyperplasia. A functional cooperation of WNT4 and TGFβ pathways in tumor growth and in its fibrotic enrichment, is not excluded. Indeed, studies showed that the TGFβ/SMAD3 pathway mobilized the WNT/β-catenin pathway in the smooth muscle cells proliferation (86) and in the development of pathological dermal fibrosis in a p38-dependent manner by decreasing the accumulation of WNT pathway negative regulators (87). In the prwKO testis, the tumor suppression is associated with a systematic reduction in the fibrotic areas that characterize the LCCSCT. Consistent with the increase of FGF2, EGF, IGF1 and KIT-L, their converging signaling pathways (MAPK and PI3K/AKT/mTOR) (88) are activated in the LCCSCTs of CNC patients and prKO mice supporting their involvement in the tumoral expansion. In addition, several studies in different tissues have shown that TGFβ signaling activates the expression of *miR-21* targeting PTEN production and activity, lifting the inhibition of the AKT/mTORC1 signaling which in turn amplifies oncogenic signal, matrix protein synthesis and fibrosis (89, 90). As revealed by the prKO transcriptomic analyses, the *miR-21* up-regulation argues in favor of such a TGFβ/mTOR crosstalk in the development of the CNC-LCCSCT. The mTOR pathway was previously shown to be activated in SC in response to *Lkb1* inactivation in a study aiming to understand the molecular mechanisms involved in testicular alterations in Peutz-Jeghers syndrome (PJS) (50). In this study, the mTOR pathway activation induced by disruption of the LKB1/AMPK pathway resulted in SC polarity and BTB alterations, resumption of their proliferative activities and loss of gem cells. However, no LCCSCT-type tumorigenesis was shown in this work, suggesting that, as in CNC-LCCSCT, the LCCSCT-PJS development could be dependent on signals activated by the mutation that should be present both in SC and other testicular compartments. In the prKO model, the PKA-induced mTORC1 pathway activation in the tumor and in SC, does not imply any alteration of the expression nor activity of AMPK, indicating that the *Prkar1a* and *Lkb1* mutations lead to mTOR pathway mobilization by a different way. In previous work examining the molecular consequences of *Prkar1a* loss in adrenal cortex leading to PPNAD lesions, we showed that PKA activity triggers an increase in TGFβ family members expression and directly activates mTORC1 to promote cell survival (91, 92). Intriguingly and by contrast with our present results in testis, we previously showed a marked reciprocal antagonism of the WNT/β-catenin and PKA pathways in adreno-cortical zonation and tumorigenesis (93) since *Prkar1a* inactivation in the adrenal cortex represses zona glomerulosa identity and opposes the pro-oncogenic action of WNT/β-catenin signaling. Altogether, our works therefore suggest that in the CNC, the molecular modalities of PKA action in PPNAD and LCCSCT expansion do not fully overlap. This would probably rely in non-identical PKA controlled key networks involved in the tissue homeostasis of adrenal and testis that yet arise from common primordium. In this context, the question of WNT-PKA cooperation in the pathogenesis of the CNC ovary remains unanswered.

In conclusion, our data demonstrate that the chronic unrepressed PKA activity in somatic testis cells of CNC patients unbalances testicular homeostasis by generating gonadal paracrine networks crisis that include an aberrant post-natal WNT4 signal acting as a driving force for shaping intratubular lesions and tumor expansion.

## METHODS

### Mice

*Sf1:Cre* (93), *Amh:Cre* (22), or *Scc:Cre* (23), *Prkar1a^fl/fl^* (9) *Wnt4^fl/fl^* (94) and *Prkaca^+/−^* mice (95) have been described previously. Mouse crossings enabled to generate multiple models: prKO (*Sf1:Cre,Prkar1a^fl/fl^* with testis somatic cell *Prkar1a* inactivation), prKO/*Prkaca^+/−^ (Sf1:Cre,Prkar1a^fl/fl^,Prkaca^+/−^* with global *Prkaca* haploinsufficiency), srKO, (*Amh:Cre,Prkar1a^fl/fl^*, Sertoli cell *Prkar1a* inactivation), lrKO, *(Scc:Cre,Prkar1a^fl/fl^*, Leydig cell *Prkar1a* inactivation), double mutant prwKO, (*Sf1:Cre,Prkar1a^fl/^*^fl^,*Wnt4^fl/fl^*) and srwKO, (*Amh:Cre,Prkar1a^fl/^*^fl^,*Wnt4^fl/fl^)*. Mouse models are enriched with *Rosa26R-mtmg* system to follow recombination events. Littermate control animals were used in all experiments. Control animals were *Prkar1a^fl/fl^* or *Prkar1a^fl/fl^*,*Wnt4^fl/fl^* depending on experiments. All mice were maintained on a mix genetic background of C57BL/6J and 129. Harvested testes were either frozen in liquid nitrogen or fixed in 4% PFA. For the BTB integrity analysis, intratesticular injection of EZ-Link Sulfo-NHS-LC-Biotin (15μl, 7.5mg/mL) was performed in the left testis (21335, Thermo Scientific). Testes were harvested 30 min after injection and fixed in 4% PFA. For embryonic analyses, the morning of vaginal plug detection was considered as E0.5. For fertility test, males were mated each night with two females for fifteen days.

### Samples from patients

Histological slides of testis lesions from 3 CNC patients were obtained from CS and HL. Human adult normal testis sections were used as control for histological analyses (UM-HuFPT151, CliniSciences). The clinical data summary of CNC patients is provided in table S1.

### Hormonal measurements

Mice were euthanized by decapitation and blood was collected between 9:00(a.m.) and 10:00(a.m.) since animals were maintained with 12h light (07:00-19:00) / dark cycle. Plasma testosterone concentrations were determined using ELISA kit for srKO/ lrKO models (MBS267437, MyBioSource) and by LC-MS/MS for prKO model (Collab. IP). Plasma LH and FSH concentrations were determined using a multiplex assay (MPTMAG-49K, Merck Millipore).

### Histology

Testes were embedded in paraffin and 5μm section were prepared. H&E, Masson’s trichrome, Red Alizarine S or Toluidine Blue stainings were performed depending on experiments. Immunohistochemistry and TUNEL staining were performed according to conditions described in table S2. Images were acquired with a Zeiss Axioplan 2, Zeiss AxioImager or Zeiss Axioscan Z1 slide scanner. A detailed description of materials and methods is provided in Supplementary material.

### Micro-array analyses

Gene expression profiles for 2-weeks and 2-month-old WT, prKO, srKO and lrKO testes were analyzed using Affymetrix Mouse Gene 2.0 ST Arrays. Raw and processed data are deposited on NCBI Gene Expression Omnibus (GEO) platform. Gene expression was normalized by RMA (Affy R package) and genotype comparisons were performed using the Limma package. All p-values were adjusted by the Benjamini-Hochberg correction method. Genes with adjusted p-value<0.05 were considered significantly deregulated and highly deregulated if abs(Log2FC)>1.0 (Datafiles S1). Heatmaps were generated with R and represent colour-coded individual median centered gene expression levels. Gene set enrichment analyses were conducted using GSEA 4.0.2 with MSigDB or custom gene sets (Datafiles S2). Permutation were set to 1000 and were performed on gene sets. Enrichment analyses of GO and KEGG terms were conducted using DAVID 6.8 (Datafiles S3). Dotplot and Volcanoplot were generated with R.

### RTqPCR

Total mRNAs were extracted using RNAII nucleotide extraction kit (Macherey Nagel) and reverse transcribed as previously described (93).Primers used are listed in table S3. For each primer pairs, efficiency of PCR reactions was evaluated by amplification of serial dilutions of a mix of cDNAs. Relative gene expression was obtained by the *ΔΔCt* method with normalization to average expression of *Actin*.

### Western blot

Thirty micrograms of total proteins were loaded on 8% SDS-page gel, transfer on nitrocellulose and detected with primary antibodies (table S2). Signals were quantified with ChemiDoc MP Imaging System camera system (Bio-rad) and Image Lab software (Bio-rad). Expression of phosphorylated or active proteins were normalized to expression of the corresponding total protein.

### Statistics

Statistical analyses were performed using R and GraphPad Prism 8. Number of mice per group are reported in the figures. All bars represent the mean ± SD. Normality of data was assessed using Shapiro-Wilk normality test. For comparison of two groups, variance equality was tested using F test. Statistical analysis of two groups was performed by two-tailed Student’s t test with or without Welch’s correction (as a function of variance) for normally distributed data or by Mann-Whitney’s test for not normally distributed data. For comparison of multiple groups, variance equality was tested using Brown-Forsythe test. Statistical analysis of multiple groups was performed by one-way ANOVA followed by Tukey multiple correction test or Welch’s one-way ANOVA followed by Games-Howell multiple correction test (as a function of variance) for normally distributed data or by Kruskal-Wallis’ test followed by Dunn’s multiple correction test for not normally distributed data. Statistical analysis of hyperplasia/tumor proportion was performed by two proportion Fisher’s test.

### Study approval

All patients undergo tumor resection and gave informed consent for the use of their resected tissues for research purposes. Mouse experiments were conducted according to French and European directives for the use and care of animals for research purposes and was approved by the local ethic committee (C2E2A).

## AUTHOR CONTRIBUTIONS

CD, AMLM and AM designed research. FG, A.Swain and SV provided *Cre* expressing and *Wnt4*-floxed mice. CAS, CK, FRF, HL and AGL provided human tumor samples and clinical data. CD, DD, AL, AMLM, NM, IP, JCP, ISB and JW performed experiments. CD, AMLM, AM, ISB and A.Septier analyzed data. CD, AMLM wrote the paper. AM, IT and PV edited the paper.

## ACKNOWLEDGMENTS

We acknowledge Dr. C.Lyssikatos (now at the University of Indiana, Indianapolis, IN, USA) and Dr. E.Belyavskaya (NICHD, NIH, Bethesda, MD, USA) for the coordination of the clinical studies and sample retrieval and shipment, under protocol 95CH0059. We thank A.François, Professor J.C.Sabourin (Head of the CRB-TMT in Rouen) and the CRB-TMT in Rouen to provide us human tumor samples. We thank K.Ouchen, S.Plantade and P.Mazuel for animal care, C.Damon-Soubeyrand (Anipath-Clermont), and J.P.Saru for their technical assistance, Y.Renaud for management of bioinformatic platform.

## Data and materials availability

The microarray data have been deposited in the Gene Expression Omnibus (GEO) Database.

## SUPPLEMENTARY METHODS

### Histology

Images were minimally processed for global levels with ZEN. Images settings and processing were identical across genotypes. Tumor scoring was performed on H&E stained sections. Testis was noted (1) either normal if any modification of stromal cell number has been observed (2) either hyperplastic if cell number is increased and (3) either tumoral if testis developed stromal nodules representing at least 20% of total section area. 25 random ST were counted to assess the percentage of ST with spermatozoa, with vacuoles, with biotin or filled of germ cells in mice; or the number of G9A+, SYCP3+, ZBTB16+ or TUNEL+ per ST in mice. Epithelium thickening and CLDN11 expression domain were measured in 25 ST per mouse. 500 stromal cells were counted to assess the PCNA proliferation index. Mast cells were detected following toluidine blue staining and were counted in a whole testis section. In human samples, the number of germ cell per tubule and the CLDN11 expression domain were quantified in 50 ST whereas 200 ST were counted to assess the percentage of ST with spermatozoa or with vacuoles.

## SUPPLEMENTARY FIGURES

**Figure S1.**
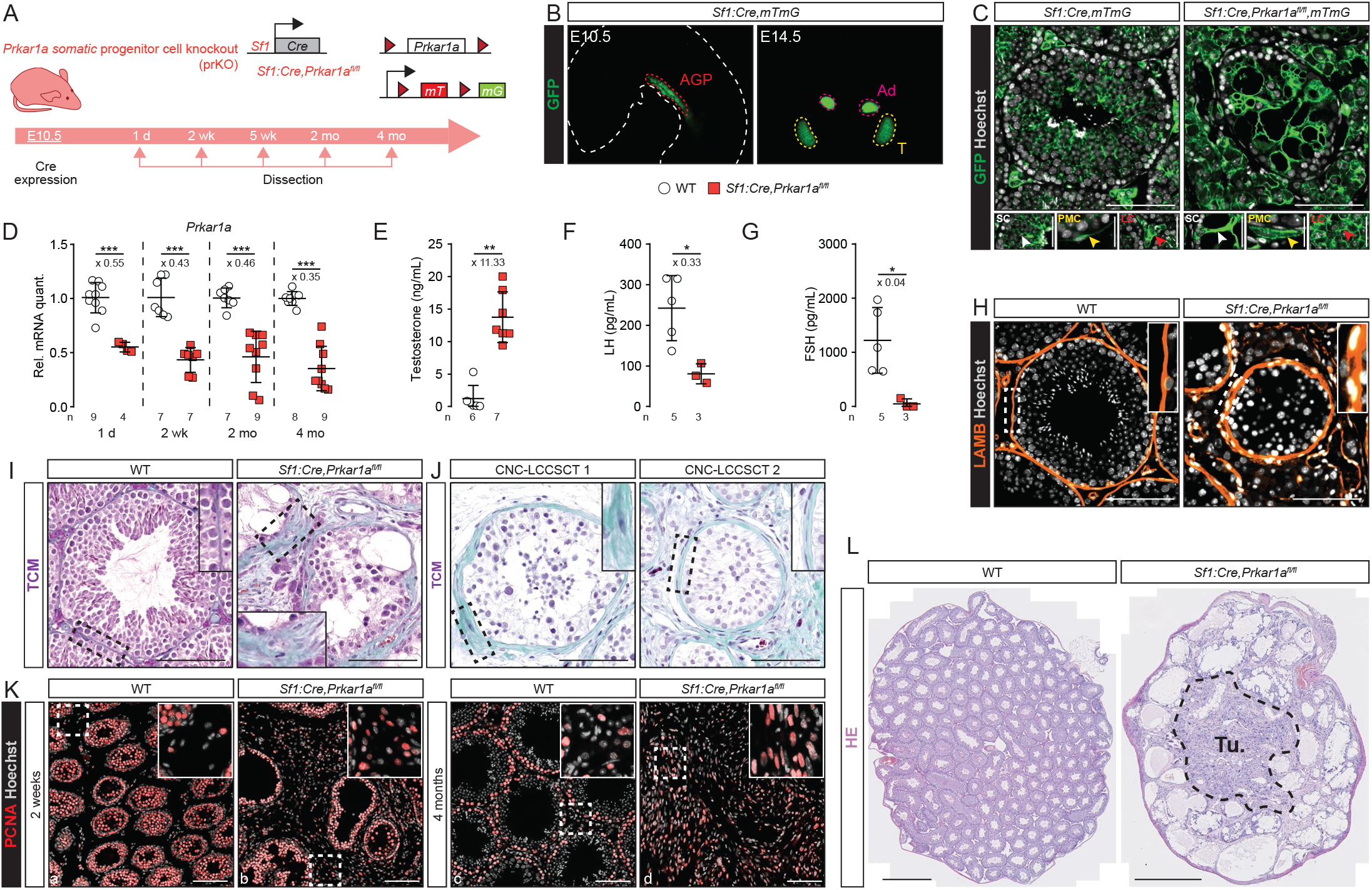
related to Figure 1. (**A** and **B**), *Prkar1a* floxed alleles were deleted in SF1+ cells. GFP is expressed from the R26R^mTmG^ locus following Sf1:Cre-mediated recombination from E10.5 in both WT (*Sf1:Cre,mTmG,Prkar1a^+/+^*) and prKO testis (*Sf1:Cre,mTmG,Prkar1a^fl/fl^*). (**C**), Immunohistochemical detection of GFP localized in SF1+ derived cells in 2-mo-old WT and prKO testes. White arrowheads, Sertoli cells (SC); yellow arrowheads, peritubular myoid cells (PMC); red arrowheads, Leydig cells (LC). Scale bars, 100μm and 50μm. (**D**), RTqPCR analysis of mRNAs levels encoding *Prkar1a* in 1-d, 2-wk, 2-mo, 4-mo-old WT and prKO testes. Statistical analysis was performed using Student’s t-test or Welch’s t-test. (**E**), Plasma testosterone concentration in 3-mo-old prKO mice compared with WT mice. (**F** and **G**), Plasma FSH and LH concentrations in 3-mo-old prKO mice compared with WT mice. Statistical analysis was performed using Mann-Whitney’s test. (**H-J**), Immunohistochemical detection of LAMB (H) and TCM staining (I-J) show peritubular thickening both in 4-mo-old prKO testis and human CNC-LCCSCT. (**K**), Immunohistochemical analysis of PCNA in 2-wk and 4-mo-old WT and prKO testes. (**L**), HE staining of central tumor in 4-mo-old prKO testis. Bars represent the mean per group ± SD. Scale bars, 100μm (inset 50μm, (L) 500μm). ns, not significant, *\*p<0.05*, *\*\*p<0.01*, *\*\*\*p<0.001.*

**Figure S2.**
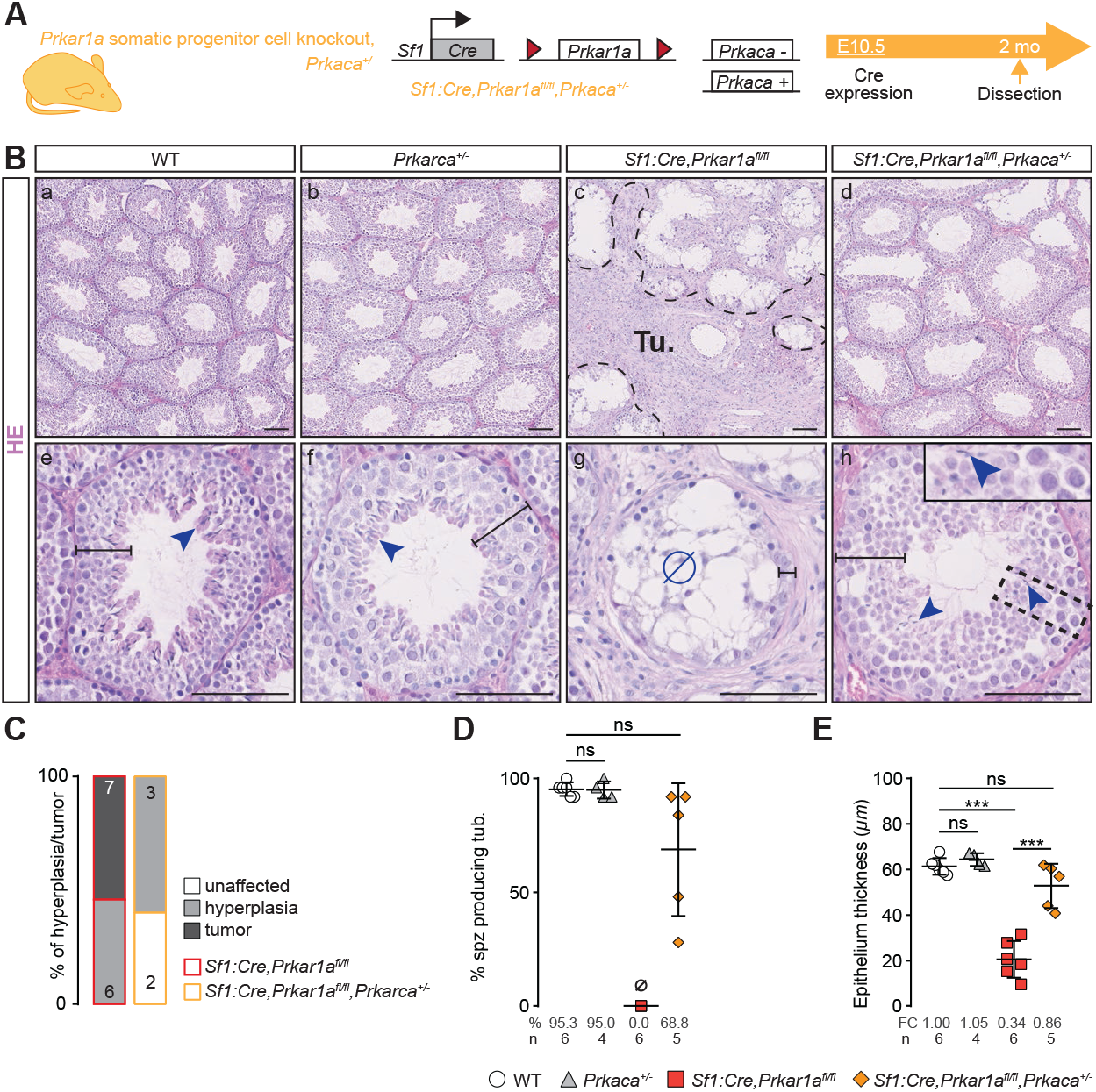
related to Figure 1. (**A**), *Prkar1a* floxed alleles were deleted in SF1+ cells and *Prkaca* heterozygosity context (noted as prKO/*Prkaca^+/−^*, *Sf1:Cre,Prkar1a^fl/fl^,Prkaca*^+/−^). (**B**), HE staining of 2-mo-old WT, *Prkaca*^+/−^, prKO and prKO/*Prkaca^+/−^* testis. Dashed black lines delineate tumor developed in prKO testis, blue arrowheads show restored spermatozoa differentiation in prKO/*Prkaca^+/−^* testis and black lines limit seminiferous epithelium thickness. Ø, absence of spermatozoa. Scale bars, 100μm. (**C**), Relative proportion of testicular hyperplasia and tumor in 2-mo-old *Prkaca*^+/−^, prKO and prKO/*Prkaca^+/−^* testes. (**D** and **E**), Percentage of ST with elongated spermatids (D) and epithelium thickness (E) quantified following HE staining in 2-mo-old WT, *Prkaca*^+/−^, prKO and prKO/*Prkaca^+/−^* testes are restored in prKO/*Prkaca*+/− testis. Bars represent the mean value per group ± SD. Statistical analysis was performed using one-way ANOVA followed by Tukey multiple correction test. *\*\*\*p<0.001.*

**Figure S3.**
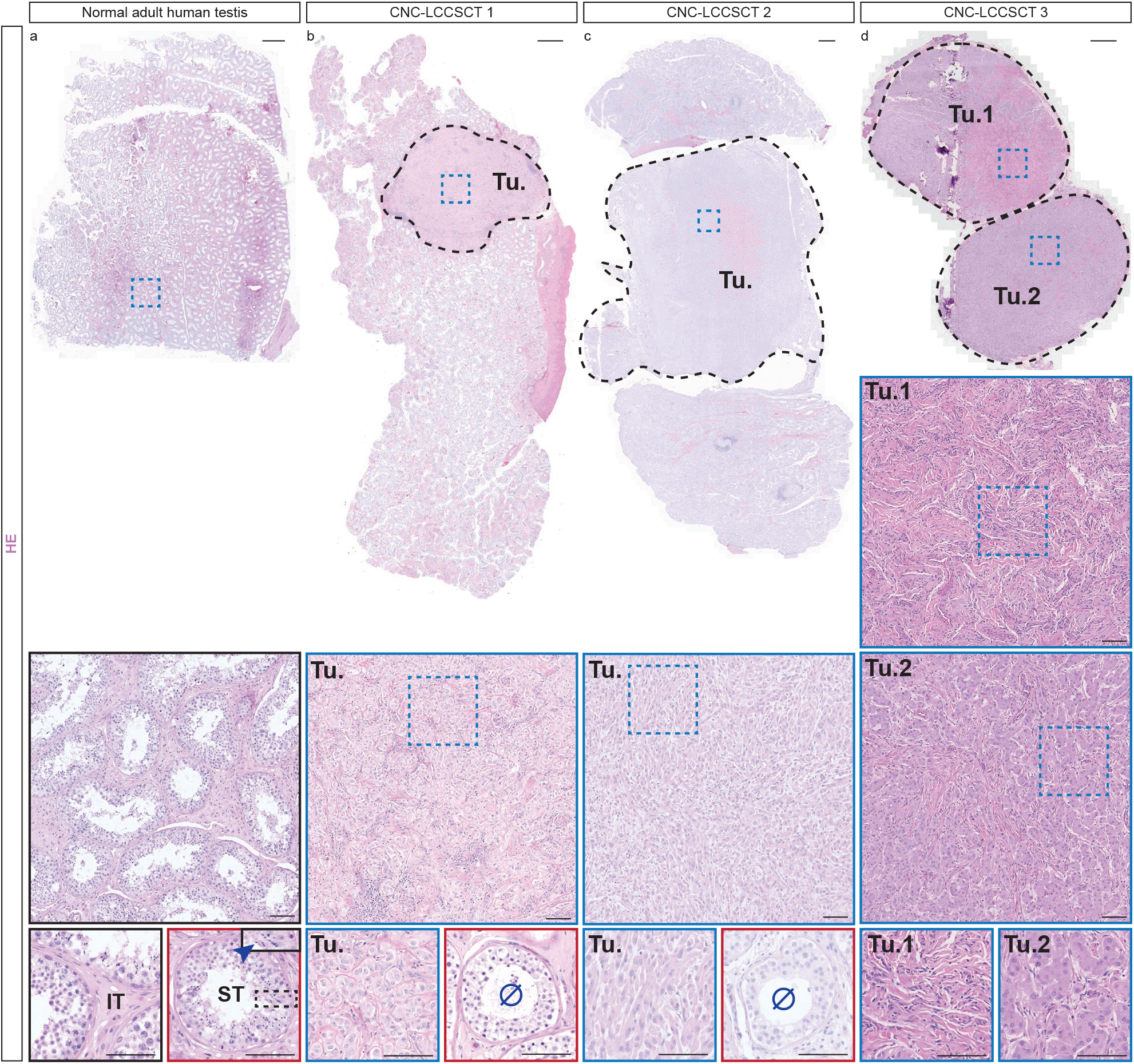
related to Figure 1. HE staining of reconstituted normal adult human testis (a) and human LCCSCT biopsy sections (b-d). Dashed lines delineate primary tumor mass. CNC-LCCSCT 3 section (d) presents two distinct tumor masses (T1, T2) and is devoid of ST. Blue square, tumor; red square, ST. Blue arrowheads, spermatozoa; Ø, disorganized ST devoid of spermatozoa. Scale bars reconstituted biopsies, 1mm; scale bars insets, 100μm.

**Figure S4.**
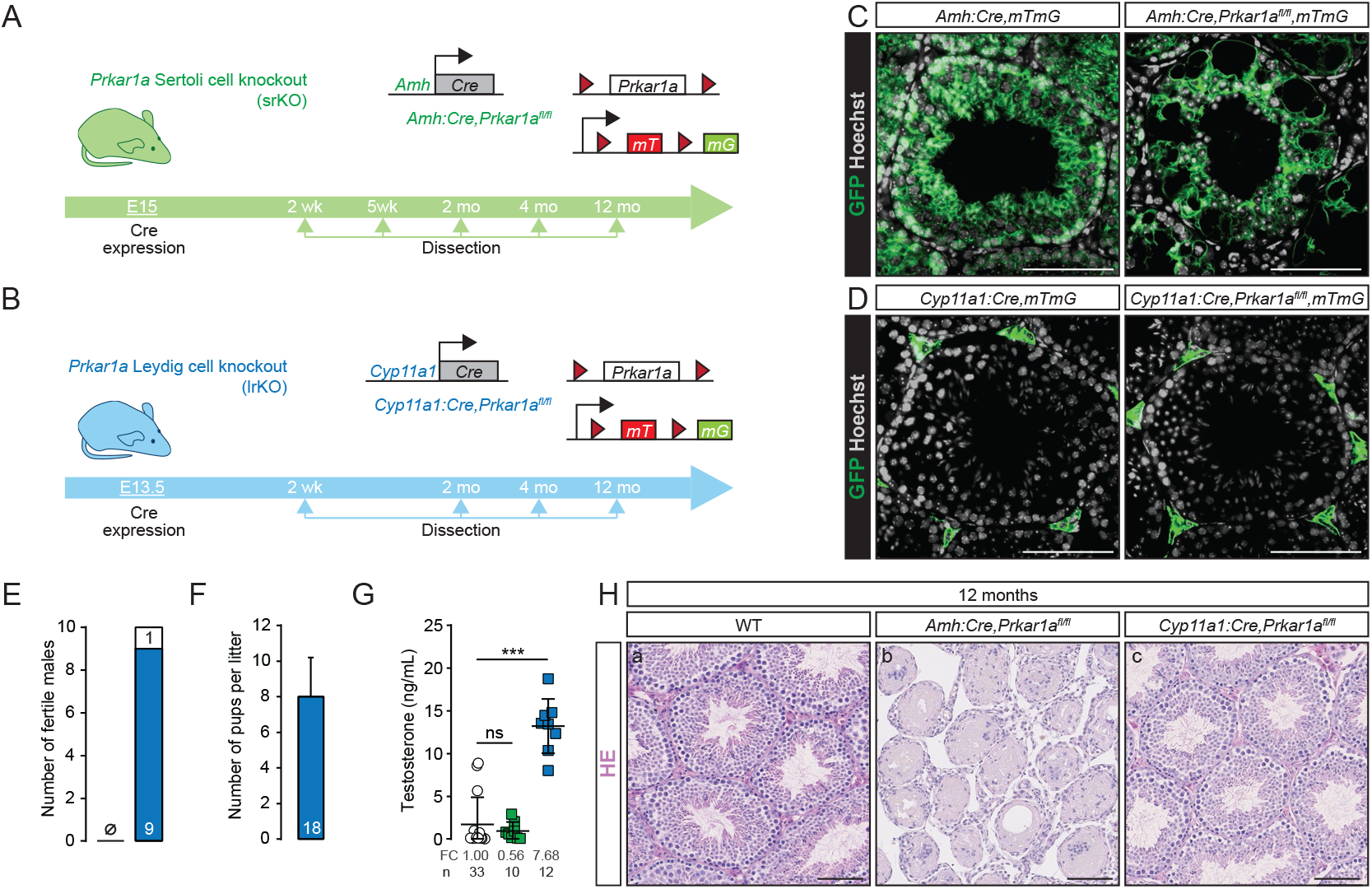
related to Figure 2. (**A** and **B**), *Prkar1a* floxed alleles were deleted in AMH+ cells (A) or CYP11A1+ cells (B). GFP is expressed from the R26R^mTmG^ locus following Amh:Cre-mediated recombination from E14.5 in srKO testis (*Amh:Cre,mTmG,Prkar1a^fl/fl^*) or following Cyp11a1:Cre-mediated recombination from E15.0 in lrKO testis (*Cyp11a1:Cre,mTmG,Prkar1a^fl/fl^*). (**C** and **D**), Immunohistochemical detection of GFP localized in AMH+ derived cells (Sertoli cells) in 2-mo-old srKO testis (C) and in CYP11A1+ derived cells (Leydig cells) in 2-mo-old lrKO testis (D). (**E** and **F**), Number of fertile males (E) and pups per litter (F) in 2-mo-old lrKO male mice. (**G**), Plasma testosterone concentrations in 2-mo-old WT, srKO and lrKO mice. Bars represent the mean per group ± SD. Statistical analysis was performed using Kruskal-Wallis’ test followed by Dunn’s multiple correction test. (**H**), HE staining of 12-mo old WT, srKO and lrKO testes showing the absence of tumor in all three genotypes. *\*\*\*p<0.001.* Scale bars, 100μm.

**Figure S5.**
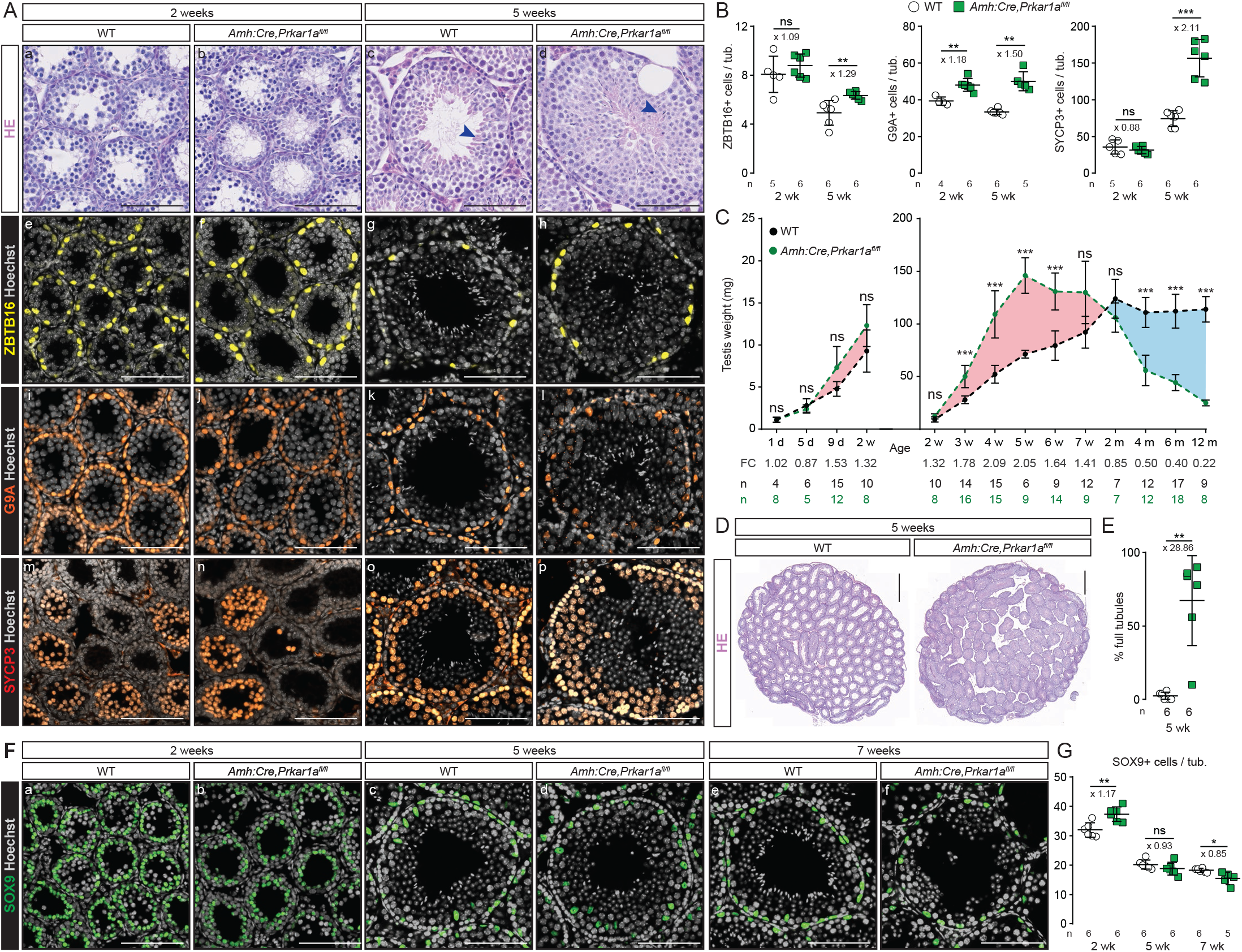
related to Figure 3. (**A**), HE staining (a-d) and immunohistochemical detection of ZBTB16 (e-h), G9A (i-l), SYCP3 (m-p) in 2-wk and 5 wk-old WT and srKO testes. Blue arrowheads, spermatozoa. (**B**), Quantification of ZBTB16, G9A and SYCP3 based on the number of positive cells per tube. Statistical analysis was performed using Student’s test or Welch’s t-test. (**C**) WT and *Prkar1a* mutant testis weight from 3-wk to 4-mo. Welch’s one-way ANOVA was followed by Games-Howell multiple correction test. (**D**), Representative HE of 5-wk-old WT and srKO sections. (**E**), Quantification of full tubules represented as a percentage of positive tubules quantified following HE staining in 5-wk-old WT and srKO testes. Statistical analysis was performed using Mann-Whitney’s test. (**F**), Immunohistochemical detection of SOX9 in 2-wk (a,b), 5-wk (c,d) and 7-wk-old (e,f) WT and srKO testes. (**G**), Quantification of SOX9 based on the number of positive cells per tube. Statistical analysis was performed using Student’s test or Welch’s t-test. *\*p<0.05*, *\*\*p<0.01*, *\*\*\*p<0.001.* Bars represent the mean per group ± SD. Scale bars, 100μm.

**Figure S6.**
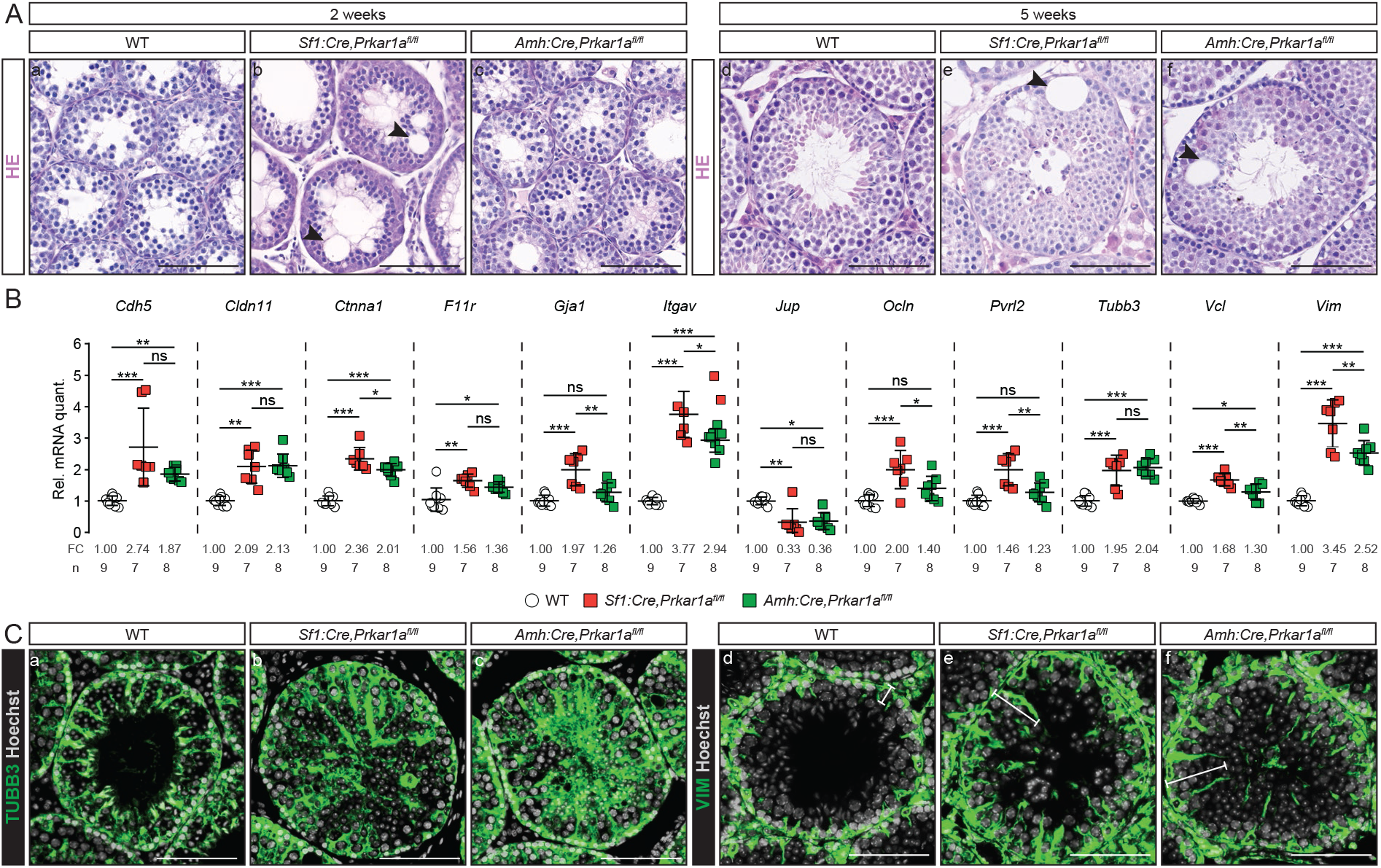
related to Figure 4. (**A**), HE staining of 2-wk (a-c) and 5-wk-old (d-f) WT, srKO and prKO testes. Arrowheads indicate vacuoles. (**B**), RTqPCR analysis of mRNAs levels encoding genes involved in cell junction assembly (*Cdh5*, *Cldn11*, *Ctnna1*, *F11r*, *Gja1*, *Itgav*, *Jup*, *Ocln*, *Pvrl2*, *Tubb3*, *Vcl*, *Vim*) in 2-mo-old WT, srKO and prKO testes. Bars represent the mean per group ± SD. Statistical analysis was performed using one-way ANOVA followed by Tukey multiple correction test, Welch’s one-way ANOVA followed by Games-Howell multiple correction test or Kruskal-Wallis’ test followed by Dunn’s multiple correction test. *\*p<0.05*, *\*\*p<0.01*, *\*\*\*p<0.001.* (**C**), Immunohistochemical detection of TUBB3 (a-c) and VIM (d-f) in 5-wk-old WT, prKO and srKO testes. White line delineates VIM expression domain.

**Figure S7.**
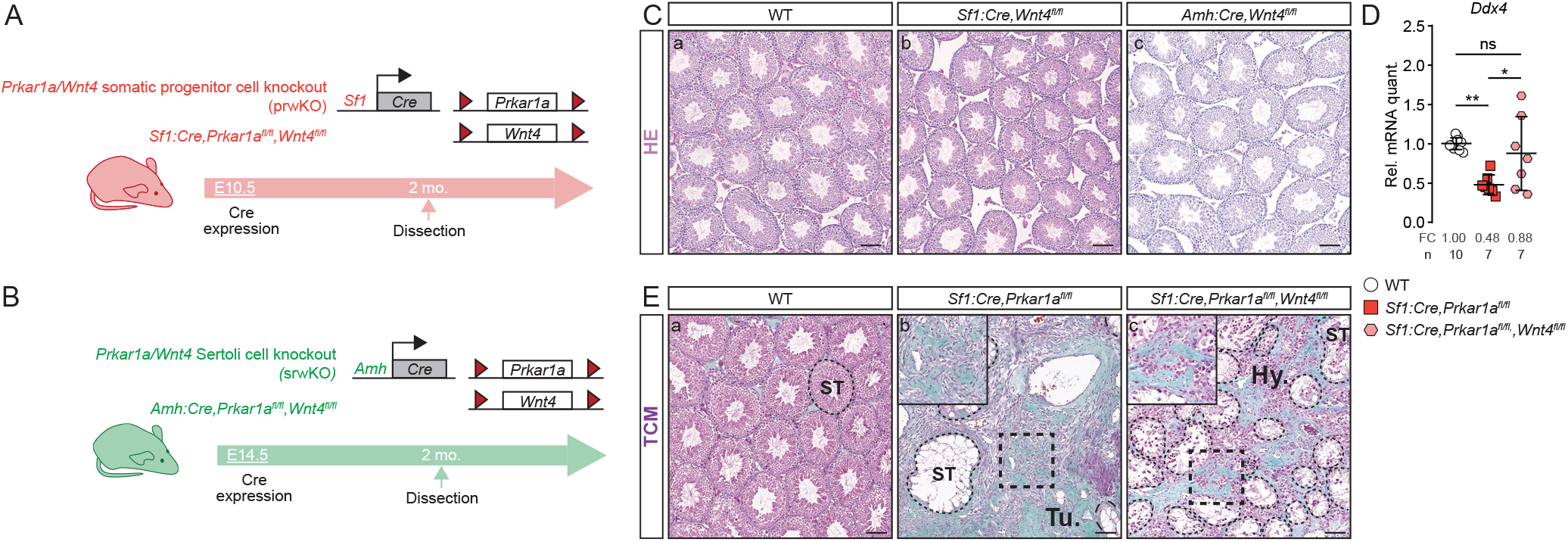
related to Figure 5. (**A** and **B**), *Prkar1a* and *Wnt4* floxed alleles were deleted in SF1+ cells (noted as prwKO, *Sf1:Cre,Prkar1a^fl/fl^,Wnt4^fl/fl^*) (A) or in AMH+ cells (noted as srwKO, *Amh:Cre,Prkar1a^fl/fl^,Wnt4^fl/fl^*) (B). (**C**), HE staining of 2-mo-old WT, *Wnt4* somatic progenitor cell knockout (noted as pwKO, *Sf1:Cre,Wnt4^fl/fl^*) and *Wnt4* Sertoli cell knockout testis (noted as swKO, *Amh:Cre,Wnt4^fl/fl^*). (**D**), RTqPCR analysis of mRNAs levels encoding *Ddx4* in 4-mo-old WT, prKO and prwKO testes. Bars represent the mean expression per group ± SD. Statistical analysis was performed using Welch’s one-way ANOVA followed by Games-Howell multiple correction test. (**E**), TCM staining of 2-mo-old WT, prKO and prwKO testes. Scale bars, 100μm. *\*p<0.05*, *\*\*p<0.01*, *\*\*\*p<0.001.*

**Table S1.** Clinical data of CNC patients

|  | CNC-LCCSCT 1 | CNC-LCCSCT 2 | CNC-LCCSCT 3 |
| --- | --- | --- | --- |
| Identification number | BH14 16096 | SI-109-208 | N518-740 |
| Associated center | Rouen, France | Bethesda, USA | Bethesda, USA |
| Carney complex | Yes | Yes | Yes |
| *Prkar1a* mutation | Yes | Yes | *PRKAR1A* genetic testing was indeterminate, due to lack of peripheral DNA testing. |
|  | ex7, c.695dup, p.Arg233LysfsX15 | c.891+3A>G p.Glu297Glu |  |
| Age | 25 years | 7 years | 6 years |
| Orchiectomy | Partial - right testis | Bilateral Orchiectomy<br>(left orchiectomy at age 7 and right orchiectomy at age 10) | Partial –left testis |
| CNC associated manifestations | LCCSCT<br>Cardiac myxomas<br>Cutaneous myxomas<br>Pituitary microadenomas<br>Thyroid nodules | LCCSCT<br>Cardiac myxomas<br>Lentigines and conjunctival pigmentation<br>PPNAD | LCCSCT<br>Pigmentary lesions |
| Hormonal manifestations | LH 4UI/l (N: 1.7-8.6)<br>FSH 3.8UI/l (N: 1.5-12.4)<br>Testosterone 3.77ng/ml (N : 3-9)<br>Estradiol 20pg/ml (N : 10-60)<br>TeBG 26nmol/l (N : 17-34)<br>Delta-4-androstenedione 2.3ng/ml (N : 0.9-2.4) | LH 0.1 U/L<br>FSH <0.1 U/L<br>Testosterone 1.14 ng/mL (Tanner II)<br>Estradiol <10 pg/mL | Not available |

**Table S2.**
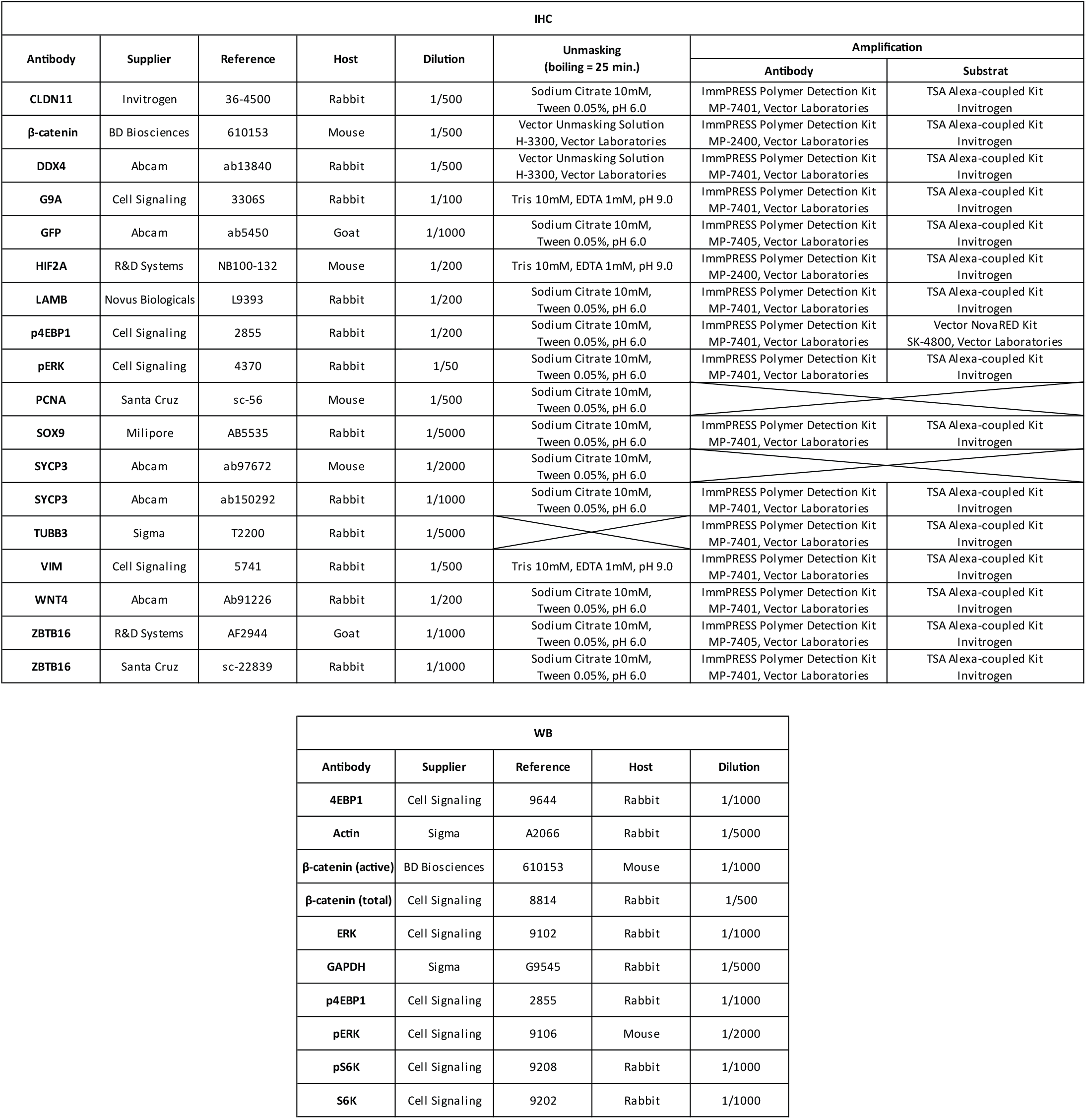
Antibodies and conditions for immunohistochemistry and western blot

**Table S3.**
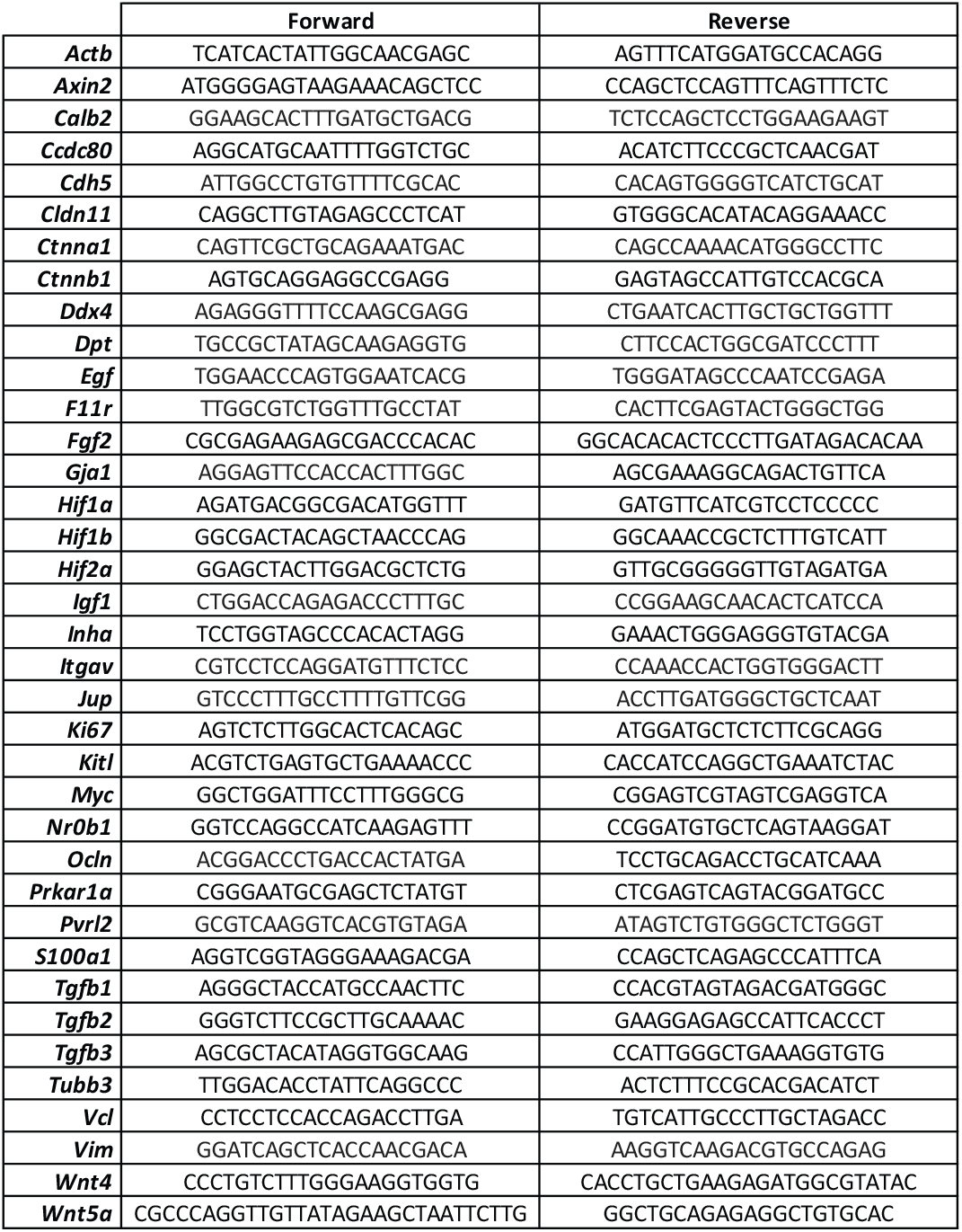
Primers used for RTqPCR

